# EagleC Explorer: A desktop application for interactively detecting and visualizing SVs and enhancer hijacking on Hi-C contact maps

**DOI:** 10.1101/2023.08.07.552228

**Authors:** Yihao Fu, Xiaotao Wang, Feng Yue

## Abstract

It has been shown that Hi-C can be used as a powerful tool to detect structural variations (SVs) and enhancer hijacking events. However, there has been no existing programs that can directly visualize and detect such events on a personal computer, which hinders the broad adaption of the technology for intuitive discovery in cancer studies. Here, we introduce the EagleC Explorer, a desktop software that is specifically designed for exploring Hi-C and other chromatin contact data in cancer genomes. EagleC Explorer has a set of unique features, including 1) conveniently visualizing global and local Hi-C data; 2) interactively detecting SVs on a Hi-C map for any user-selected region on screen within seconds, using a deep-learning model; 3) reconstructing local Hi-C map surrounding user-provided SVs and generating publication-quality figures; 4) detecting enhancer hijacking events for any user-suggested regions on screen. In addition, EagleC Explorer can also incorporate other genomic tracks such as RNA-Seq or ChIP-Seq to facilitate scientists for integrative data analysis and making novel discoveries.

## Introduction

Structural variations (SVs), including deletions, inversions, duplications, and translocations, can contribute to tumorigenesis by inducing the formation of oncogenic fusion genes and enhancer hijacking among multiple other mechanisms^1-3^. To detect such SVs in patients’ genome, scientists have been using different sequencing technologies, such as whole-genome sequencing (WGS), long-read sequencing by Nanopore or PacBio, and optical mapping^4^.

Recently, several groups including ours showed that Hi-C, a technique that was invented for the study of 3D genome folding ^5, 6^, could serve as a powerful tool to detect a full range of SVs^7-11^. The Hi-C based SV detection takes advantage of the observation that SVs induce aberrant off-diagonal or inter-chromosomal squares on a Hi-C map, and we have shown that Hi-C could uniquely detect a set of SVs that are missed by WGS and long-read sequencing^11^. However, all current software is designed to run in the Linux environment. Existing Hi-C visualization systems, such as juicebox^12^ and Higlass^13^, are designed to visualize Hi-C maps and cannot be used to predict SVs in the genome. Therefore, there is a critical need to develop software that can empower the researchers who do not have access to a Linux server or the expertise to run the pipelines, to explore such datasets, especially in a clinical setting.

Another exciting area of the 3D genome study in cancer is the enhancer hijacking model^3, 14-18^. We previously developed the NeoLoopFinder^19^, which can facilitate the discovery of gene fusions and enhancer-hijacking events genome-wide^19^. It works by reconstructing the local Hi-C maps surrounding the SV breakpoints in the cancer genome and performing loop-calling in the cancer-specific local Hi-C maps. But still, NeoLoopFinder has been designed as a Python package that requires users to have a Linux server and have programming skills, which could be a barrier for its application in the broader scientific community.

Therefore, we develop EagleC Explorer, a desktop application that can effectively detect, visualize, and examine SVs and enhancer hijacking events in real time. It works with both Mac and Windows. EagleC Explorer comes with two separate but inherently connected modes, SV Discovery mode and SV Reconstruction mode. The SV Discovery mode allows users to smoothly zoom in and out of the Hi-C maps, and navigate to any local regions via either gene names or genomic coordinates. The SV detection engine is based on our recently developed deep-learning framework EagleC^11^, so that users can easily choose a region on Hi-C with mouse and run the SV prediction. The prediction is usually finished within seconds. To visualize or predict enhancer hijacking events, users will use the SV Reconstruction mode, which reconstructs sample-specific local genome assemblies according to the types and orientations of SV breakpoints. In this mode, EagleC Explorer can plot Hi-C maps along with genes, epigenomic tracks, and 2D features on these corrected local assemblies. Like the SV Discovery mode, SV Reconstruction also allows users to zoom in and out interactively, so that users can investigate 3D genome organization changes induced by a specific SV at different scales. Switching between the two modes is seamless. Once users identify an SV breakpoint using SV Discovery, they can immediately switch to SV Reconstruction to quickly check whether this SV induces a fusion gene or novel chromatin interactions, and then they can switch back to SV Discovery mode to identify other putative SVs in the same dataset. Users can also use SVs detected by other methods and techniques as input to SV Reconstruction to examine the impact of those SVs on 3D genome, and thus intuitively prioritize SVs.

## Detection and visualization of SVs

To illustrate the utility of EagleC Explorer in detecting and visualizing SVs and enhancer hijacking, we focused on an in situ Hi-C dataset for LNCaP^19^, a prostate adenocarcinoma cell line. The Hi-C reads were mapped to the reference genome and binned at multiple resolutions (from 5kb to 5mb) as a .mcool file. We loaded the Hi-C map into EagleC Explorer under the SV Discovery mode and upon initiation, the genome-wide map was plotted on the screen. We could immediately observe the abnormally strong inter-chromosomal interactions between chr4 and chr6, chr6 and chr10, chr7 and chr14, chr6 and chr16, and chr1 and chr15 (Supplementary Figure 1), indicating the existence of inter-chromosomal translocations. Double-clicking on any particular pixel on the genome-wide view will zoom into the local region centered at that pixel. We can also zoom in a region by dragging the mouse cursor over the desired region and clicking the “zoom in” button in the pop-up dialog.

To predict SVs in a selected region, a user can simply click on the “call SV” button in the same dialog window. This would invoke the built-in EagleC module, a deep-learning model we previously developed for SV detection using chromatin interaction data^11^. If the module finds any SVs in this region, the precise SV breakpoints and the direction will be displayed on screen. In this example, the EagleC module identified an inter-chromosomal translocation between chr7:14,165,000 and chr14:37,520,000 (right lower panel in Figure 1a). Typically, candidate SV coordinates are identified and returned within seconds for any local regions. In addition to inter-chromosomal translocations, EagleC Explorer can be used to discover different types of intra-chromosomal SVs, including both long-range (>1Mb) and short-range (<1Mb) SVs (Supplementary Figures 2-4), thanks to its underlying deep-learning models^11^.

**Figure 1.**
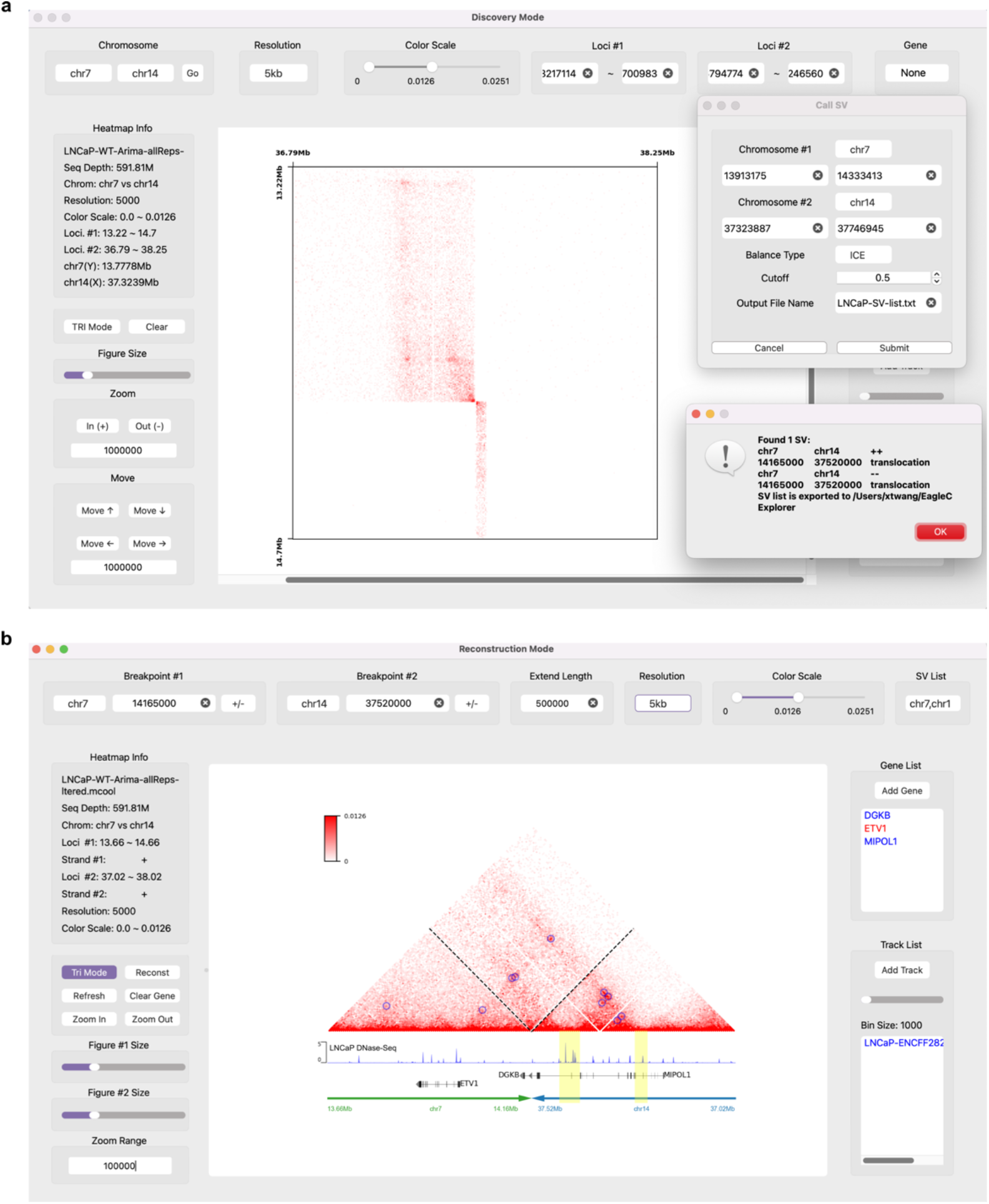
EagleC Explorer enables interactive detection of SVs and enhancer hijacking on contact maps. **a.** A screenshot of EagleC Explorer under the SV Discovery mode showing local Hi-C contacts at 5kb resolution between chr7 and chr14 in the LNCaP cell line. The toolbar at the top allows users to quickly navigate Hi-C maps between different chromosomes, resolutions, and regions. The panel on the left shows basic information about the current view. And the zoom and move buttons allow users to move and zoom in and out of Hi-C maps with configurable step sizes. In the pop-up windows, we identified one reciprocal translocation (chr7: 14,165,000; chr14: 37,520,000; ++/--) in this region. **b**. A screenshot of EagleC Explorer under the SV Reconstruction mode. The Hi-C heatmap, DNase-Seq track, and associated genes are reconstructed for the translocation event “chr7: 14,165,000, +; chr14: 37,520,000, +” in LNCaP cells. Blue circles indicate the detected chromatin loops by NeoLoopFinder. The panel on the right allows users to add gene and BigWig tracks below the heatmap.

### Detection and visualization of enhancer hijacking

Once an SV breakpoint is identified, users can go back to the genome-wide view or chromosome-wide view to identify SVs in other regions, or alternatively switch to the SV Reconstruction mode to closely check the SV just identified, without exiting the application. After selecting an SV from the dropdown menu of the loaded SV list, a Hi-C map surrounding the breakpoints will be drawn, with the breakpoint coordinates highlighted in black circles (Supplementary Figure 5). By default, the map will be shown in a “square” format. By clicking on the “Tri Mode” button on the left panel, the same map will be plotted in “triangle” format, so that other tracks, such as gene annotation and epigenomic signals, can be better accommodated and aligned with the Hi-C data. After clicking on the “Reconst” button, EagleC Explorer will reconstruct a local genome with SV fragments placed in the correct order and orientation, and plot Hi-C maps and other tracks based on it. The reconstructed map will be plotted in a separate window, so that users can compare the differences between maps before and after reconstruction. Alternatively, users can slide out any of them to just focus on one map.

Enhancer hijacking is typically achieved through the formation of neo-loops across the breakpoints. To facilitate the detection of enhancer hijacking, we integrated the Peakachu module, a machine-learning framework we previously developed for predicting chromatin loops from chromatin interaction data^20^. Activating this module is as simple as clicking on “Call Loop” from the “Functions” drop-down menu. By default, the identified chromatin loops will be displayed as blue circles on the Hi-C map. In Figure 1b, we reconstructed the local Hi-C map for the translocation described above (chr7: 14,165,000, +; chr14: 37,520,000, +), and three neo-loops were identified, with each linking the *ETV1* gene promoter with potential enhancers on chromosome 14 (indicated by DNase-Seq peaks within yellow bands), which suggests enhancer hijacking.

SV Reconstruction mode also allows users to zoom in and out of Hi-C maps with configurable step sizes (Supplementary Tutorial), and Hi-C heatmaps and all the data tracks before and after reconstruction will be synchronized by location and zoom level automatically. When adding the gene track, users can select which genes to plot from the drop-down gene list, in which genes are sorted by the genomic distance to breakpoints and cancer-related genes are highlighted in red. With all the features and modules incorporated in the package, EagleC Explorer can serve as a useful tool to view fusion genes and genes that might be influenced by neo-loops^19^.

### Summary

We developed EagleC Explorer, which is a standalone desktop application (for both Mac and Windows) for SV and enhancer detection. The graphical interface is friendly and intuitive, and all the operations illustrated above can be simply done by moving, dragging, or clicking the mouse. EagleC Explorer can directly generate publication-quality figures in various image formats such as portable network graphics (PNG), portable document file (PDF), and scalable vector graphics (SVG). We anticipate that EagleC Explorer will serve as an important tool for users to fully utilize Hi-C and other chromatin contact data to detect, visualize, examine, and prioritize SVs and enhancer hijacking events, enabling hypothesis generation about diseases and discovery of therapeutic targets.

## Data Availability

EagleC Explorer, a test dataset, and a video tutorial can be downloaded from http://3dgenome.fsm.northwestern.edu/eagleC/explorer/. Detailed tutorial is provided in the supplementary material.

## Acknowledgements

This work was supported by the National Institutes of Health [grants R35GM124820, R01HG011207, and R01HG009906 to F.Y.].

## Author Contributions

F.Y. conceived, designed, and supervised the project. Y.F. and X.W. designed and implemented the software. Y.F., X.W. and F.Y. wrote the manuscript.

## Supplementary Information

### Supplementary Figures

**Supplementary Figure 1.**
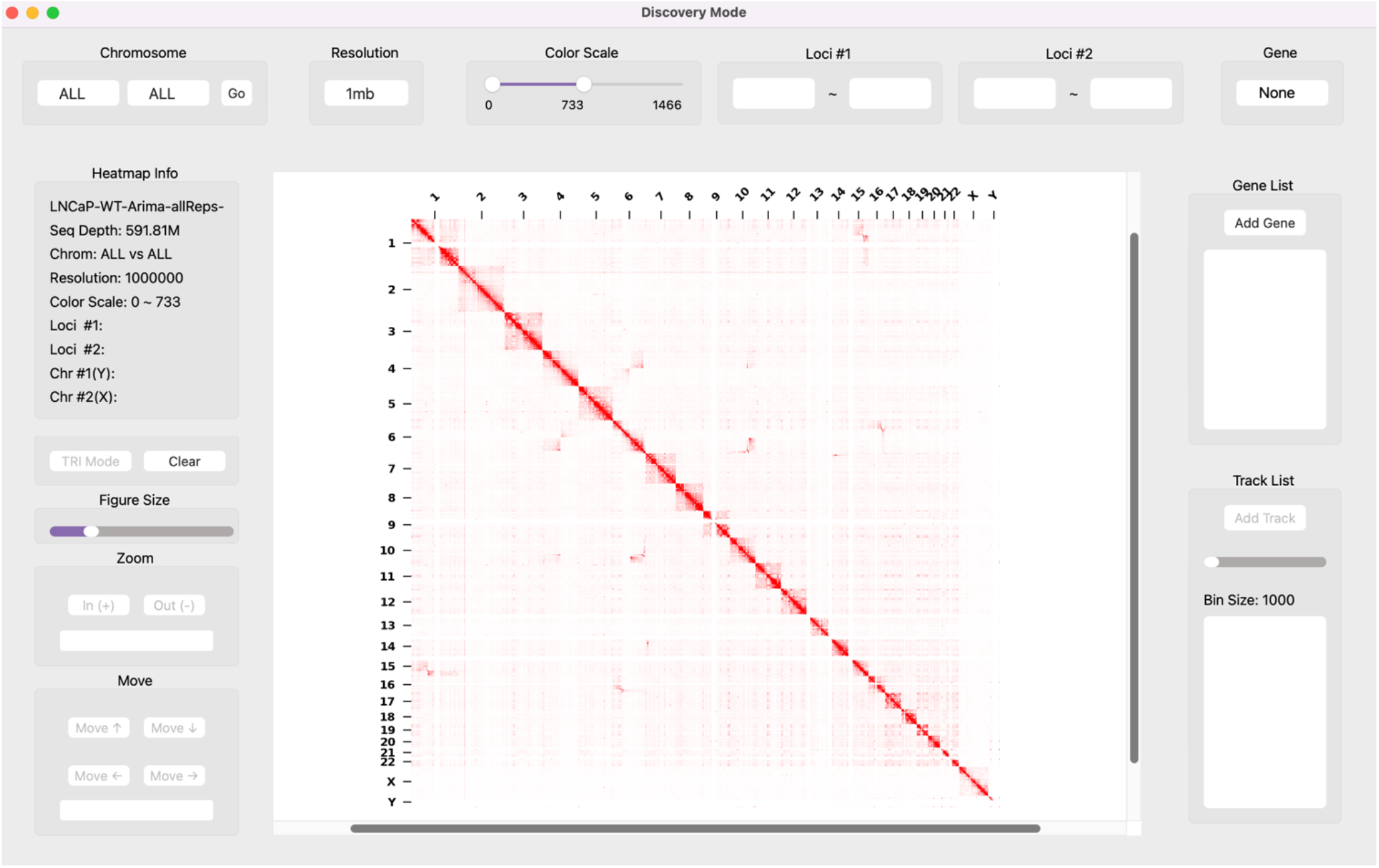
Genome-wide view of the LNCaP Hi-C map.

**Supplementary Figure 2.**
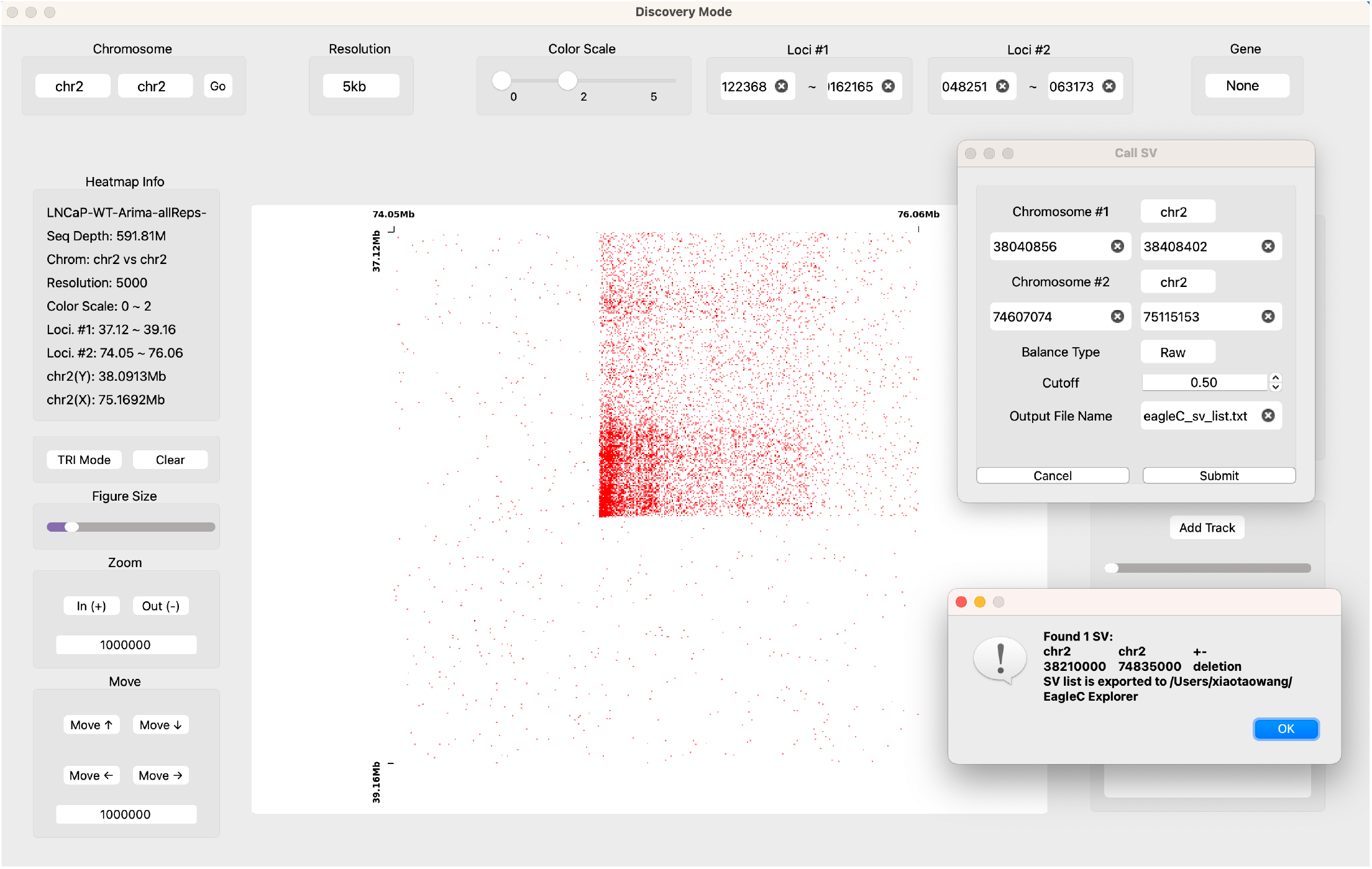
A long-range intra-chromosomal SV (chr2: 38,210,000,+; chr2: 74,835,000,-) detected by EagleC Explorer in the LNCaP cell line.

**Supplementary Figure 3.**
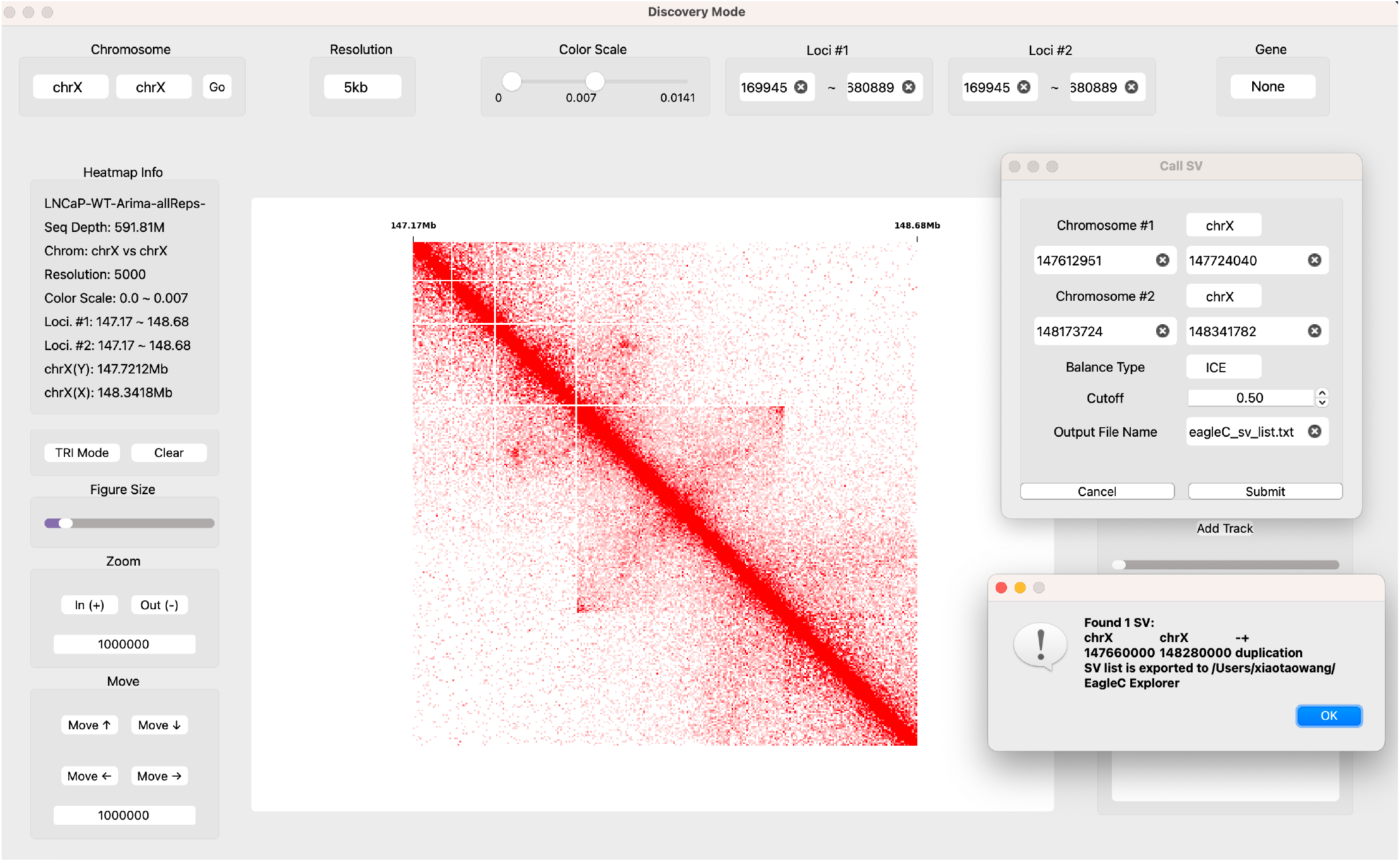
A 620-kb duplication (chrX: 147,660,000 – 148,280,000) detected by EagleC Explorer in the LNCaP cell line.

**Supplementary Figure 4.**
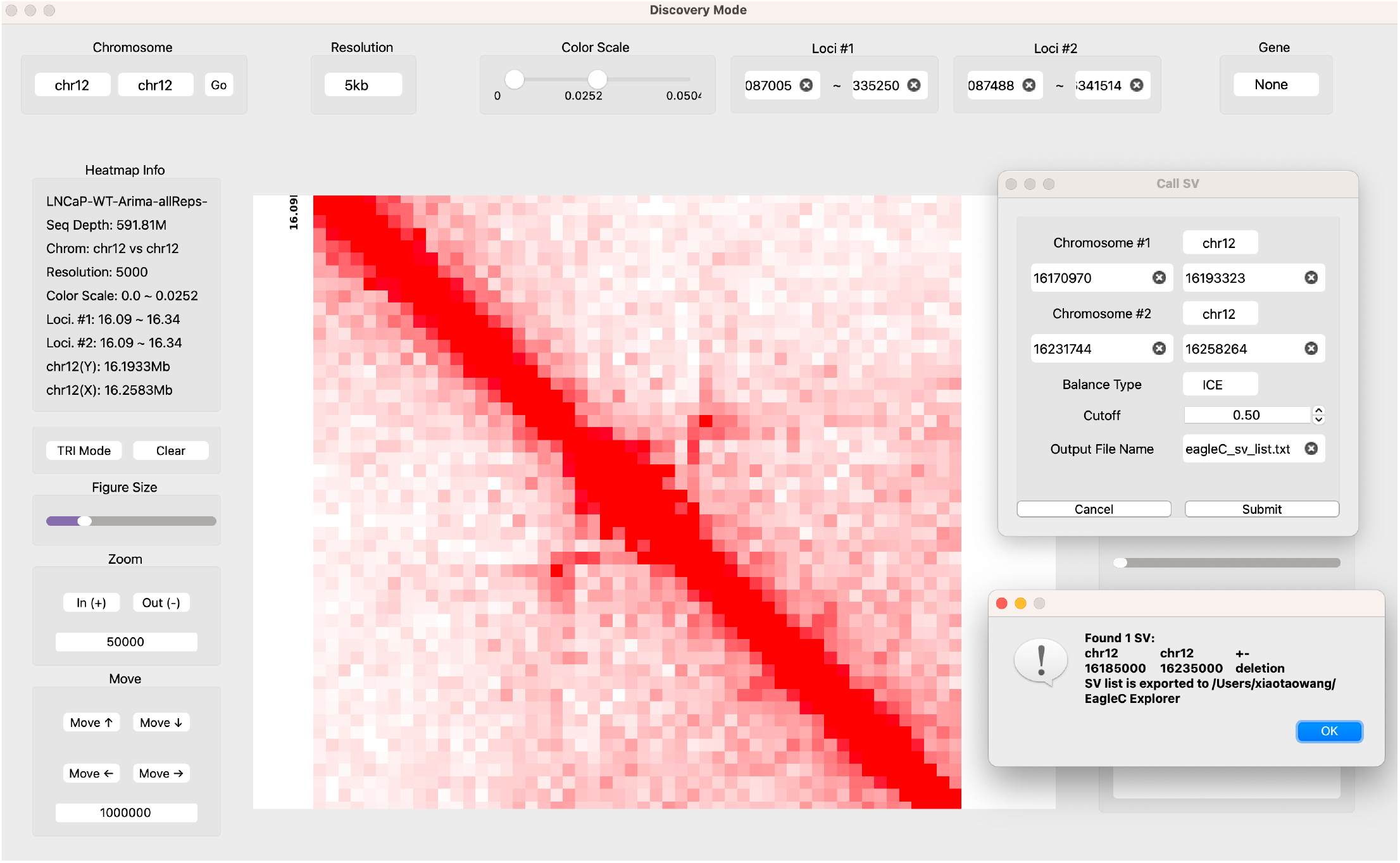
A 50-kb heterozygous deletion (chr12: 16,185,000 – 16,235,000) detected by EagleC Explorer in the LNCaP cell line.

**Supplementary Figure 5.**
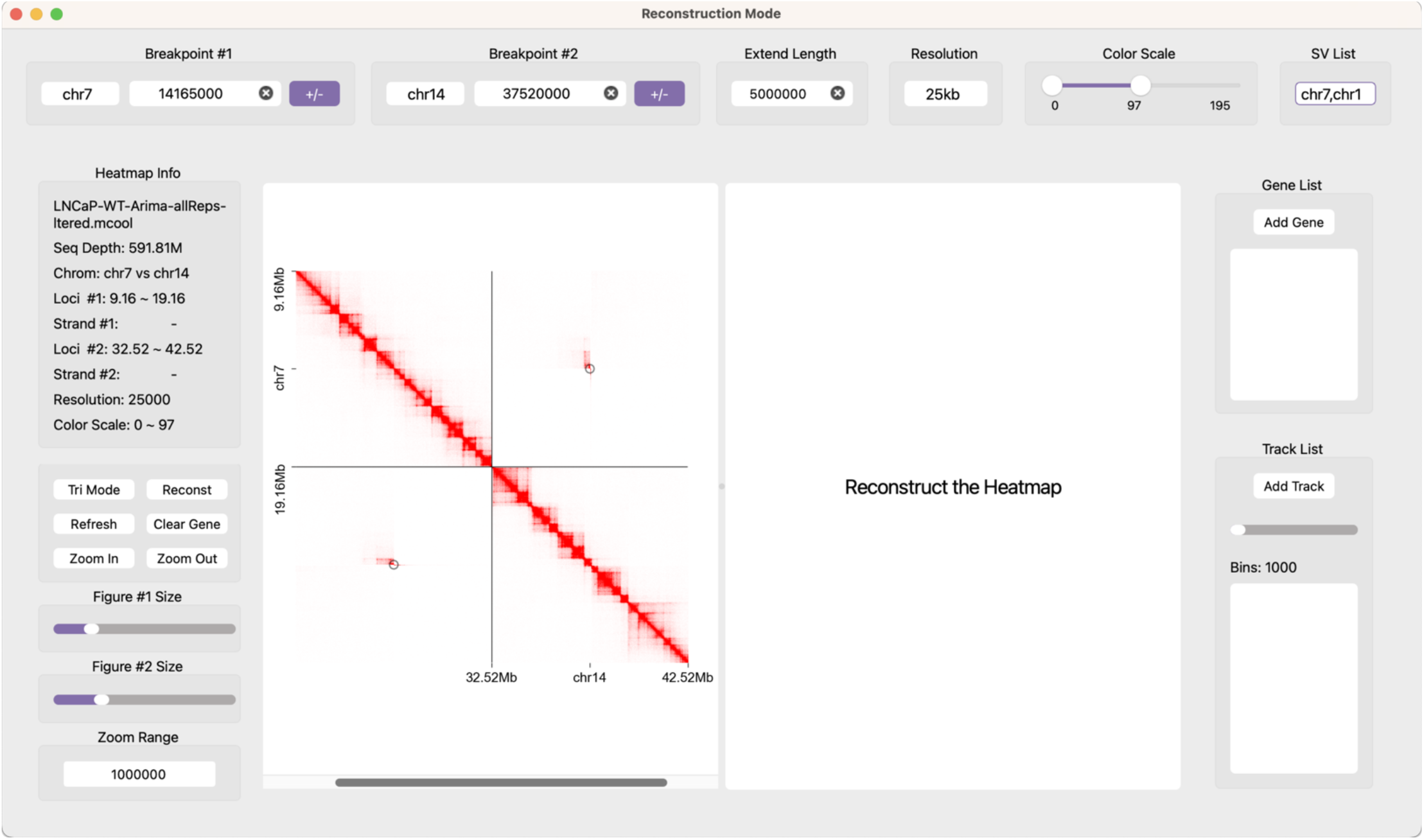
A screenshot showing the Hi-C map surrounding the breakpoints of the selected SV. The breakpoint coordinates are highlighted in black circles.

### Supplementary Methods

#### 1 Hi-C data pre-processing

For Hi-C and other chromatin contact data, EagleC Explorer accepts the cooler (.mcool) format (https://cooler.readthedocs.io/en/latest/schema.html#multi-resolution), which stores chromatin contacts at multiple resolutions from the same dataset^1^. The 4D Nucleome (4DN) Data Portal hosts “.mcool” files for >600 public Hi-C experiments, some of which are from cancer samples, serving as a great resource to explore^2^. To process Hi-C data from raw sequencing reads, we suggest using the runHiC Python package (https://pypi.org/project/runHiC/), which is an easy-to-use command-line tool based on the 4DN Hi-C data processing pipeline. Processed Hi-C data in other formats can also be converted into the “.mcool” format using existing tools, such as hic2cool (https://github.com/4dn-dcic/hic2cool) and HiCLift (https://github.com/XiaoTaoWang/HiCLift)^3^. EagleC Explorer is able to display Hi-C maps in raw, ICE-normalized, or CNV-normalized values. However, EagleC Explorer does not calculate bias vectors required for the normalization. It simply extracts bias vectors *B_i_* from either the “weight” column (for ICE normalization) or the “sweight” column (for CNV normalization) existing in a “.cool” file, and performs the transformation using the following equation:

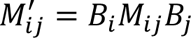

where *M_ij_* represents the raw chromatin contact frequency between bin *i* and bin *j*. The “weight” column can be calculated using the cooler Python package through command “cooler balance” (https://cooler.readthedocs.io/en/latest/cli.html#cooler-balance), and the “sweight” column can be calculated using NeoLoopFinder through command “correct-cnv” (https://github.com/XiaoTaoWang/NeoLoopFinder#cnv-normalization)^4^.

#### 2 SV format

For the SVs used in the SV Reconstruction mode, EagleC Explorer uses a 6-column tab-separated TXT format (https://github.com/XiaoTaoWang/NeoLoopFinder#format-of-the-input-sv-list). Each line in the file corresponds to the information of one SV. Sequentially, the six required fields are:

1. Chromosome of the first breakpoint
2. Chromosome of the second breakpoint
3. Orientation type of the SV. Possible values are “+-”, “+-”, “-+”, or “--”.
4. Chromosome position of the first breakpoint
5. Chromosome position of the second breakpoint
6. SV type. Values can be “deletion”, “inversion”, “duplication”, “translocation”, etc.

#### 3 Identification of chromatin loops

For chromatin loops, EagleC Explorer accepts a regular BEDPE format (https://bedtools.readthedocs.io/en/latest/content/general-usage.html#bedpe-format), and only the first six columns are required. Many methods have been proposed for genome-wide identification of chromatin loops^5,6^, and the output of most methods can be used as an input to SV Discovery mode directly or after minor format conversion efforts. For SV Reconstruction, we suggest using our recently developed toolkit NeoLoopFinder^4^, which is able to identify chromatin loops surrounding SV breakpoints, including loops across the breakpoints.

#### 4 Interactive SV detection using EagleC models

The interactive SV detection made by EagleC Explorer is based on our previously published EagleC models^7^. In brief, because different types of SVs are characterized by unique patterns on Hi-C maps, we re-framed SV detection as a classical image classification problem, and trained a series of deep-learning models to process Hi-C maps. When EagleC Explorer performs predictions, the 21 × 21 submatrices surrounding every pixel in a selected region are extracted, normalized and denoised using the same way used by EagleC, and then passed as input to the pre-trained EagleC models. For each pixel, EagleC Explorer calculates a probability score for each orientation (+-, +-, -+, and --), and a pixel will be returned as a candidate SV breakpoint if the maximum probability score exceeds the predefined cutoff which can be changed in the pop-up “call SV” dialog.

### Supplementary Tutorial

#### 1 Introduction

EagleC Explorer includes two modes (**Tutorial Figure 1**), SV Discovery and SV Reconstruction. SV Discovery allows users to view and detect SVs interactively on regular Hi-C maps, while SV Reconstruction facilitates the visualization of Hi-C maps along with genes, one-dimensional (1D) genomic tracks, and two-dimensional (2D) features on the reconstructed SV local assembly.

When EagleC Explorer is started for the first time, it will download the pre-trained EagleC models and gene database automatically, and save these files into a folder named “EagleC Explorer” in your home directory.

**Tutorial Figure 1.**
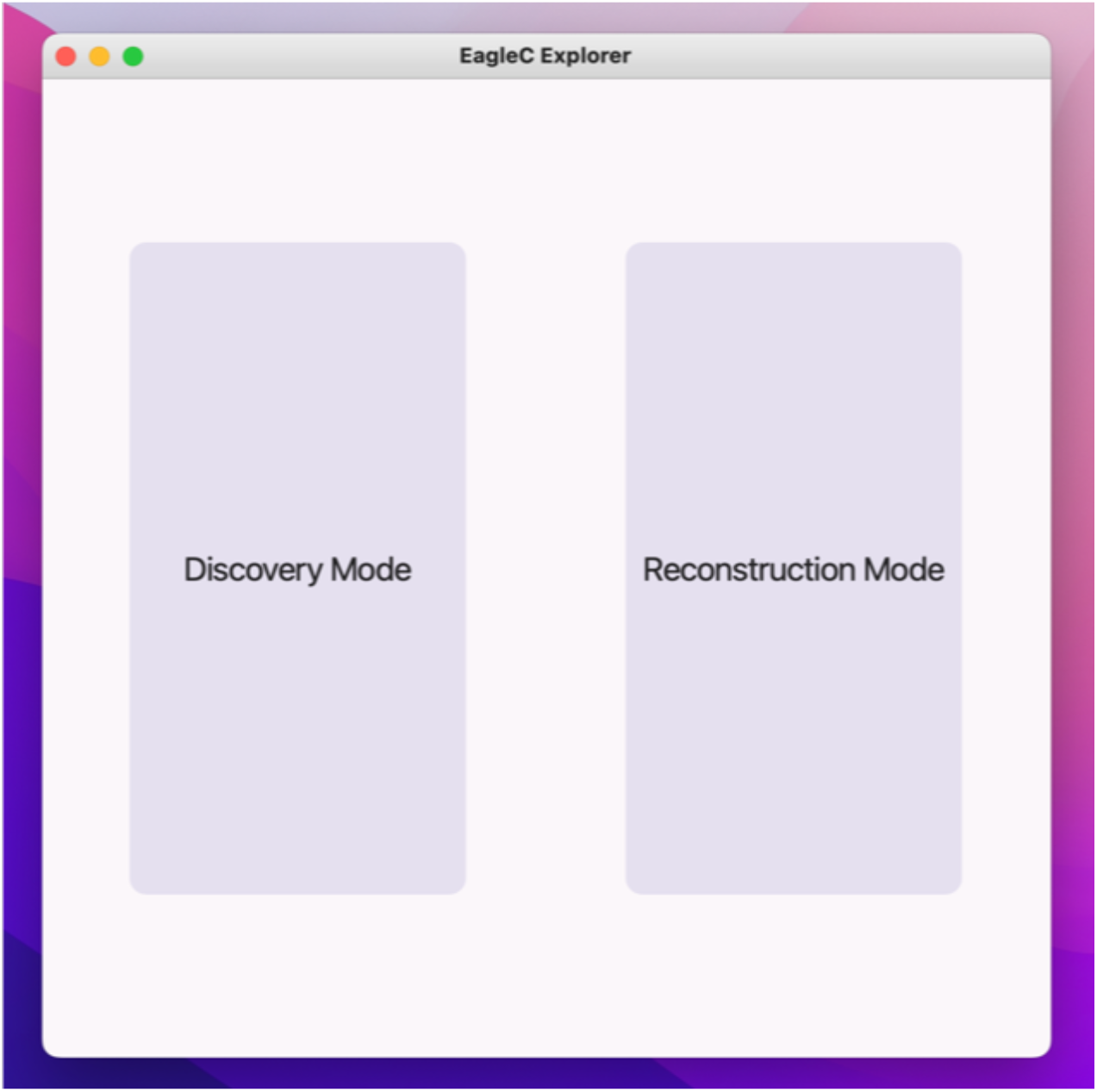

#### 2 The SV Discovery Mode

##### 2.1 Import Data

SV Discovery allows user to visualize a regular Hi-C map in the multi-resolution cooler format (.mcool) (refer to this link for detailed descriptions of this format: https://cooler.readthedocs.io/en/latest/schema.html#multi-resolution). To load/import “.mcool” files, user can either click “Import Cooler” (shortcut “command + I”) from the “File” menu (**Tutorial Figure 2**), or drag a “mcool” file into the center of the application (**Tutorial Figure 3**). Once a “.mcool” file is loaded, a genome-wide view of the Hi-C map will be displayed at the 1Mb resolution by default. Detailed information about the current view, including file name, sequencing depth of the Hi-C data, chromosome name, resolution, color scale, and genomic coordinates of current region, will be displayed on the left panel (**Tutorial Figure 4**).

**Tutorial Figure 2.**
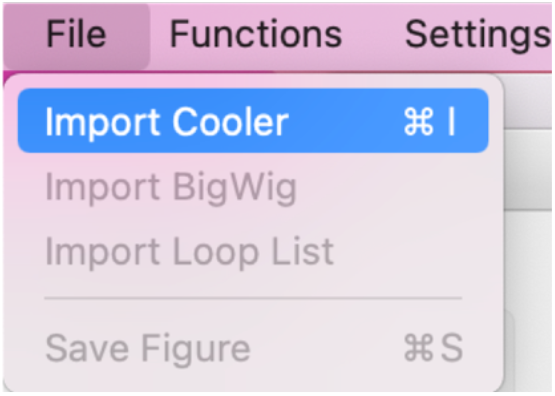

**Tutorial Figure 3.**
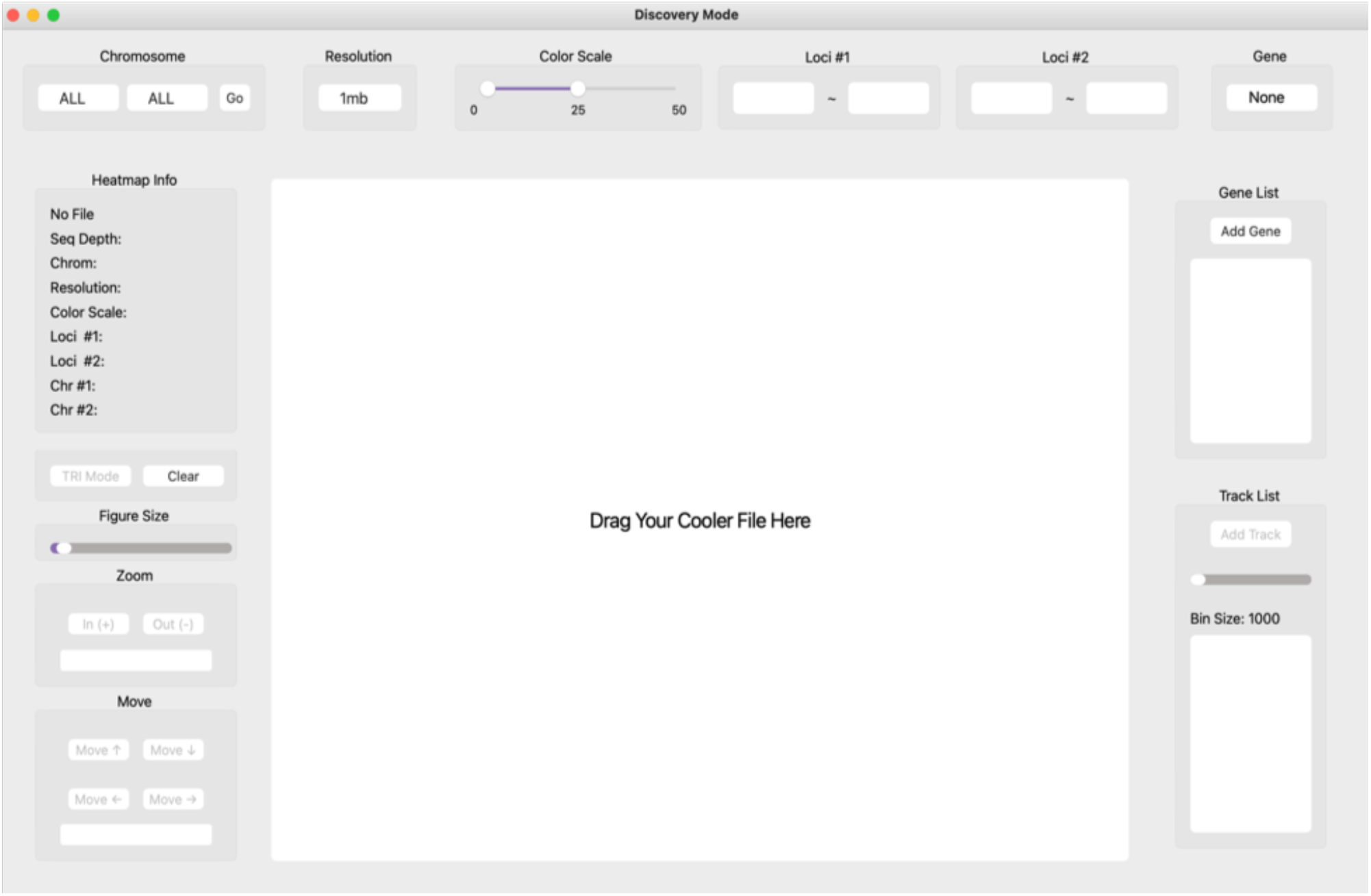

**Tutorial Figure 4.**
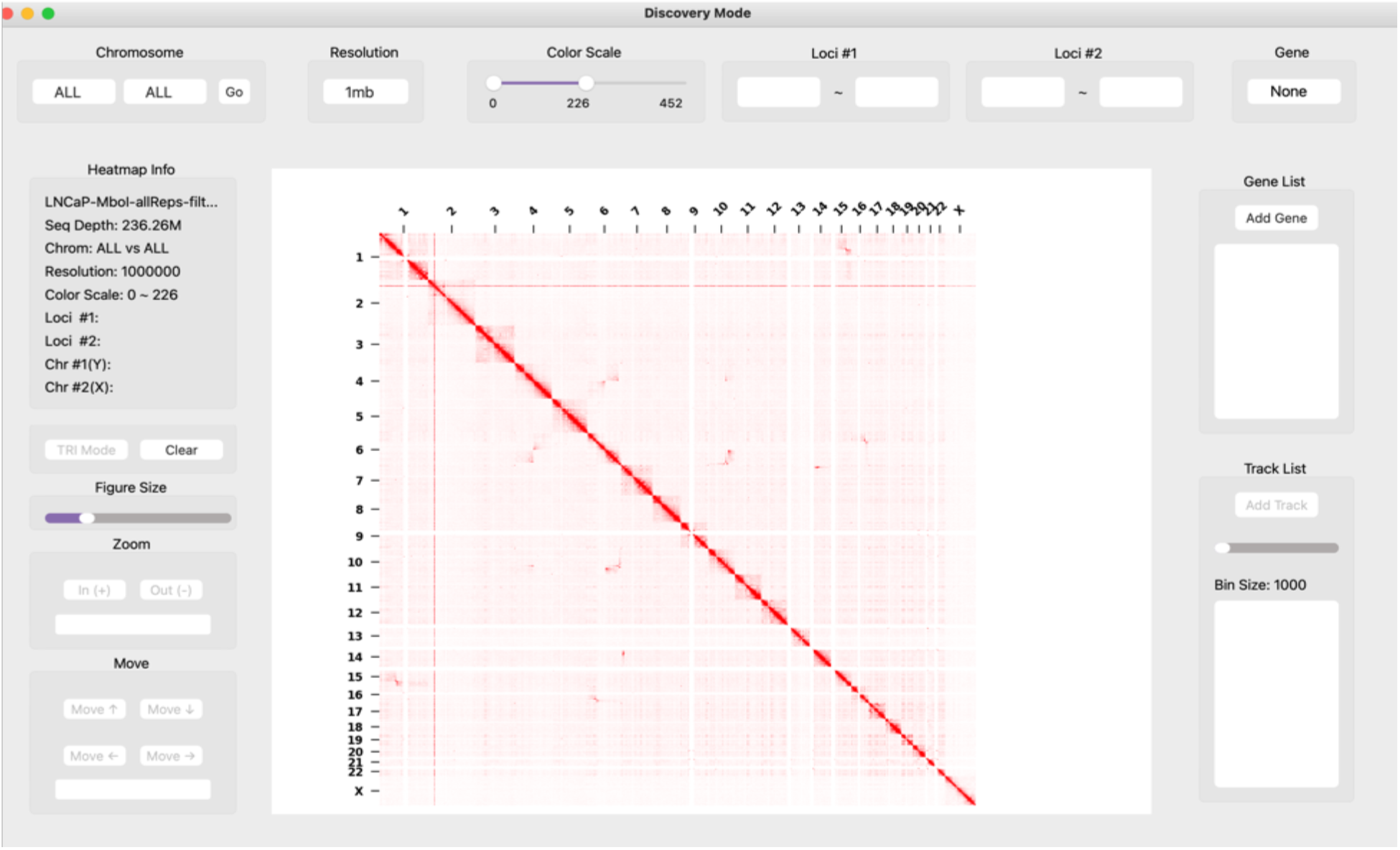

##### 2.2 Navigate Hi-C maps between different chromosomes, resolutions, and regions

By using the drop-down chromosome menu on the top-left corner, users can quickly check Hi-C maps between any two chromosomes (**Tutorial Figure 5**). By inputting valid genomic intervals (in base pairs) in the “Loci #1” and “Loci #2” text boxes and clicking the “go” button, users can further navigate the Hi-C map to a local region on the specified chromosomes (**Tutorial Figure 6**). Clicking the “Clear” button will make the current view back to the chromosome-wide map. And clicking “Clear Loci Input” in the “Functions” menu will clear all input text in the “Loci #1” and “Loci #2” boxes.

**Tutorial Figure 5.**
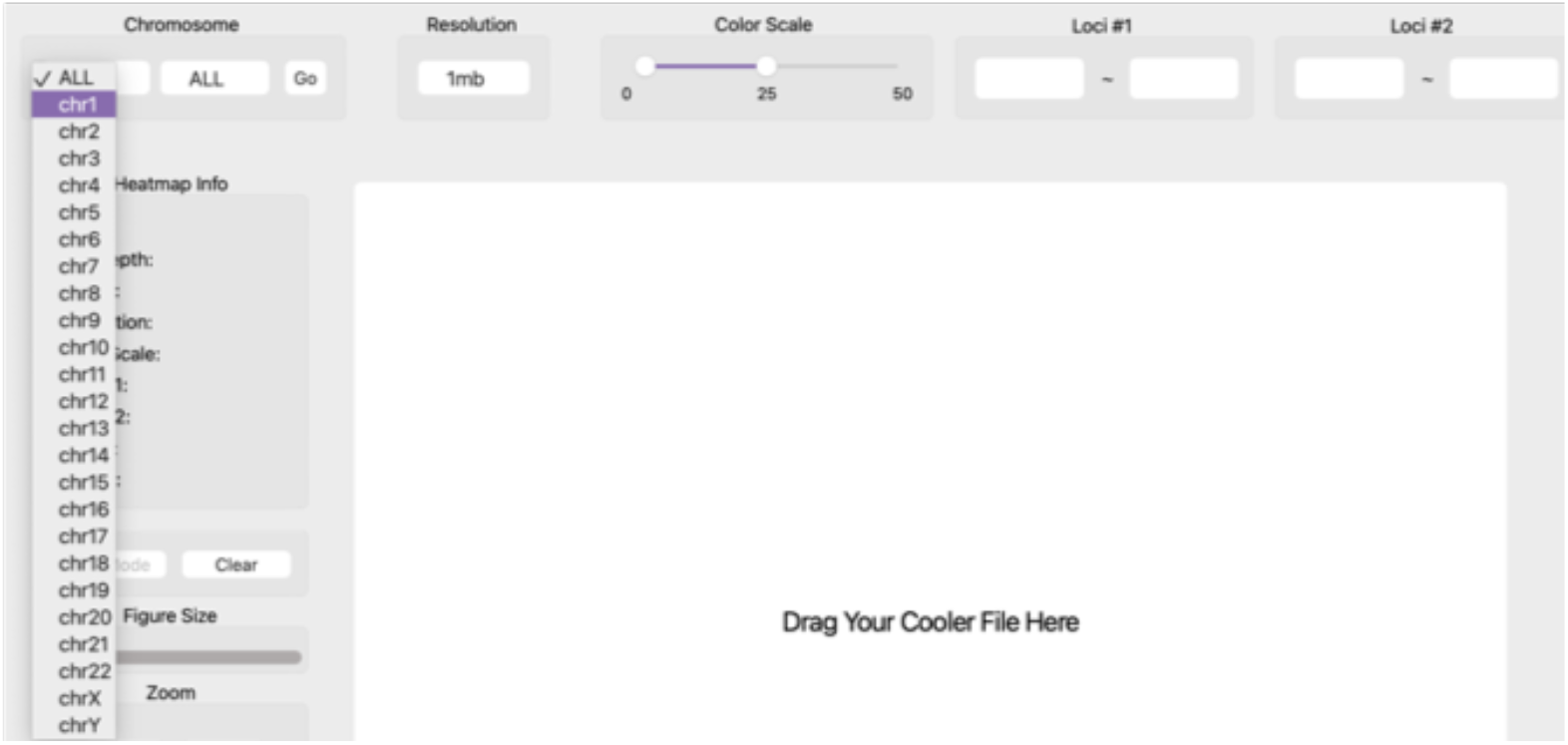

**Tutorial Figure 6.**
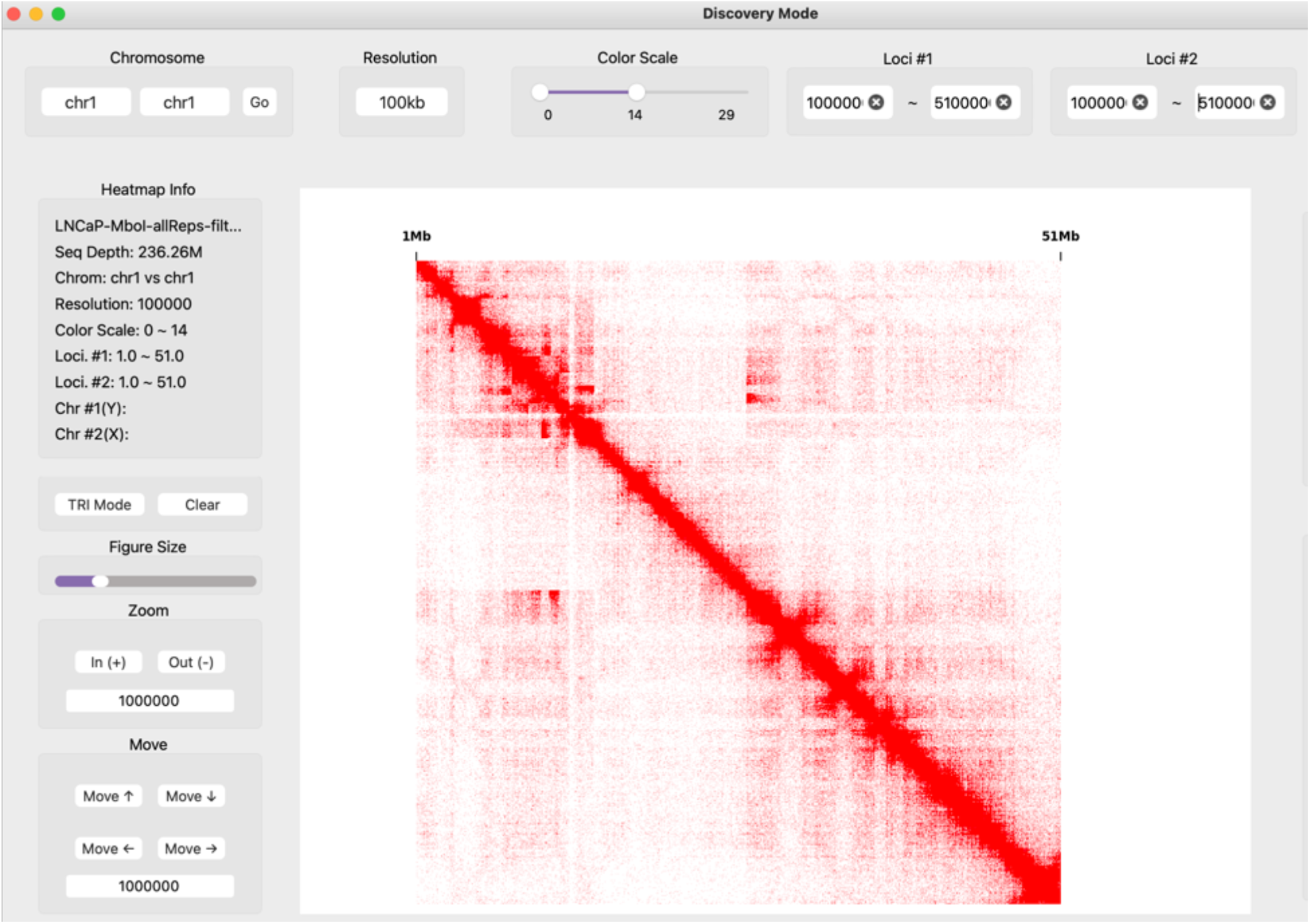

##### 2.3 Change the color scale

By default, the maximum value covered by the heatmap is set to the 95 percentiles for intra-chromosomal maps, and to the 100 percentiles for inter-chromosomal maps. This value can be adjusted through “Color Scale Percentile” in the “Settings” menu (**Tutorial Figures 7-8**).

**Tutorial Figure 7.**
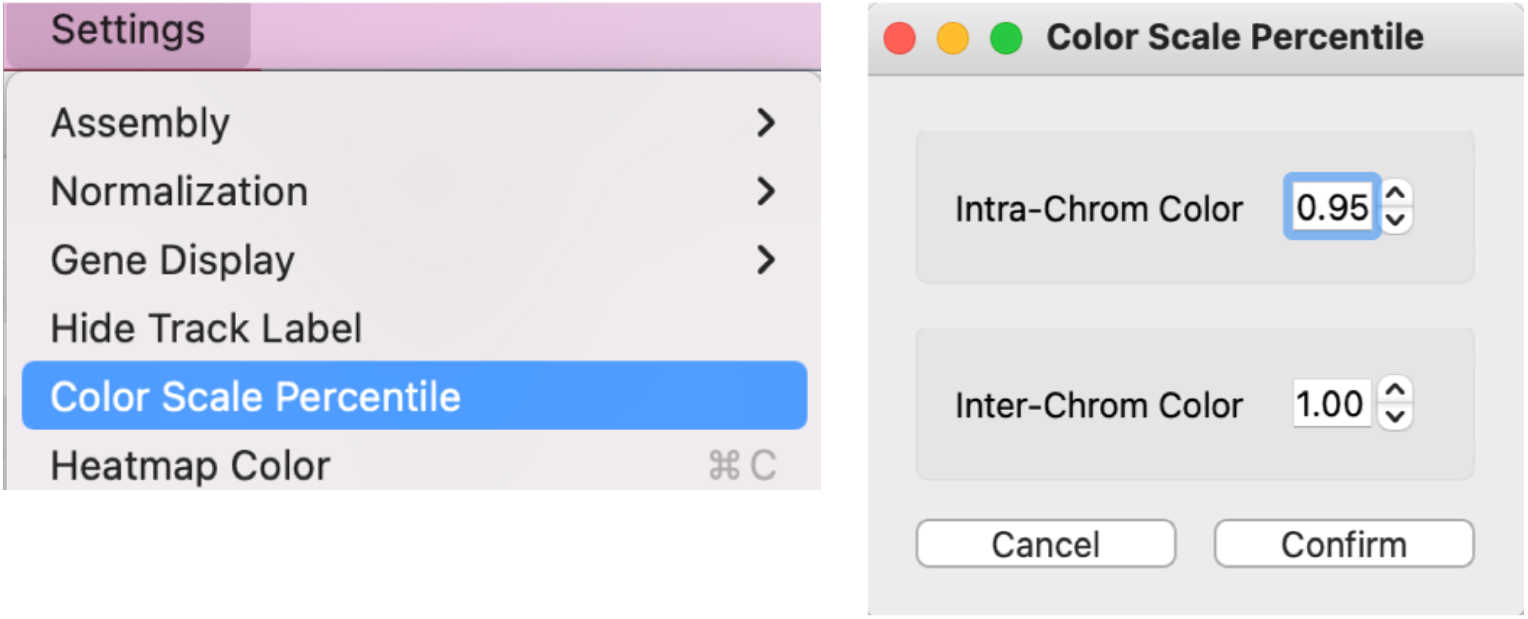

**Tutorial Figure 8.**
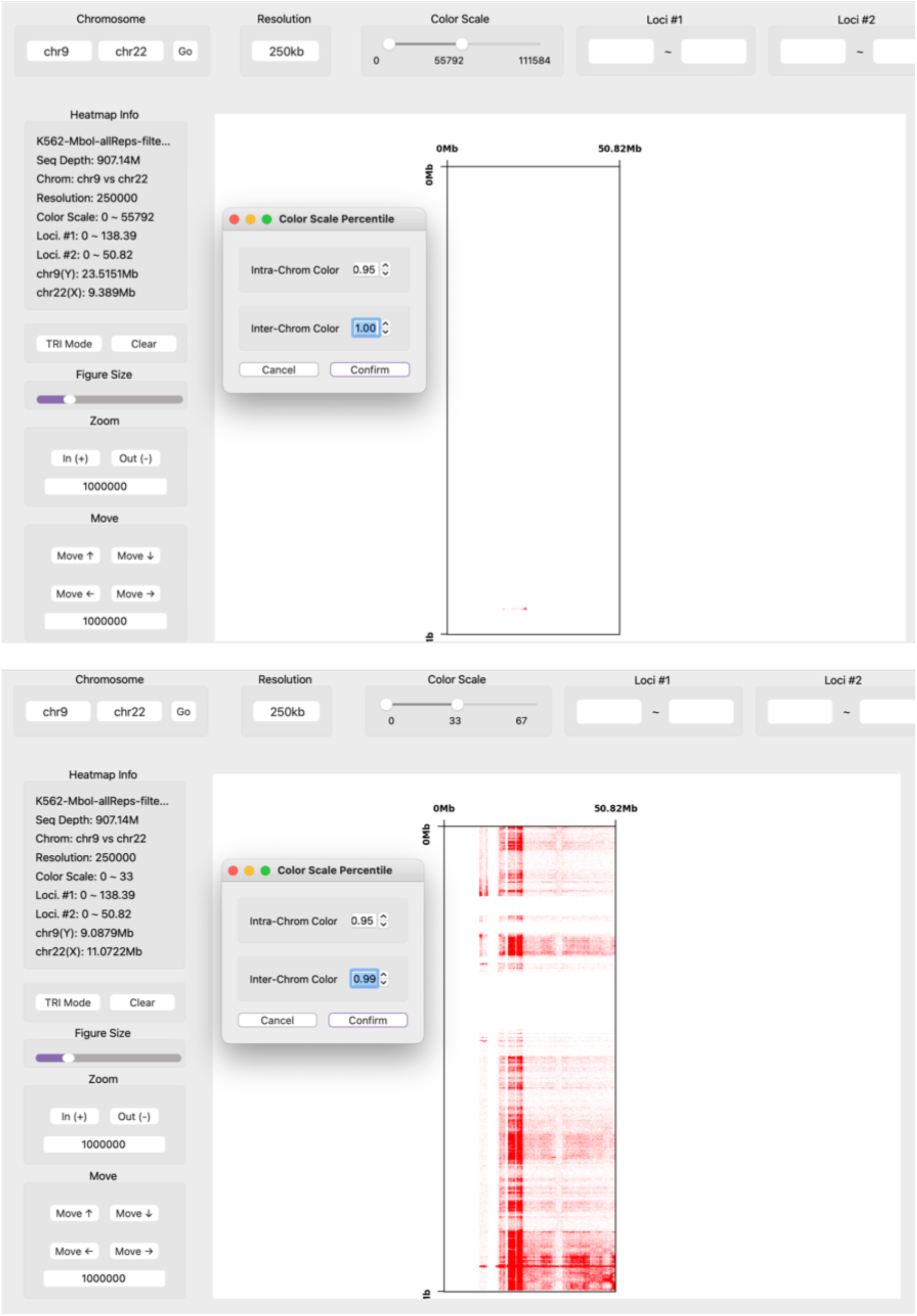

##### 2.4 Plot Hi-C maps in a “triangle” format

Clicking the “TRI Mode” button below the information panel will display Hi-C maps in a “triangle” format (**Tutorial Figure 9**). And clicking on the same button when the “triangle” format is displayed will switch back to the “square” format. Note that the “triangle” format is only available for intra-chromosomal maps.

**Tutorial Figure 9.**
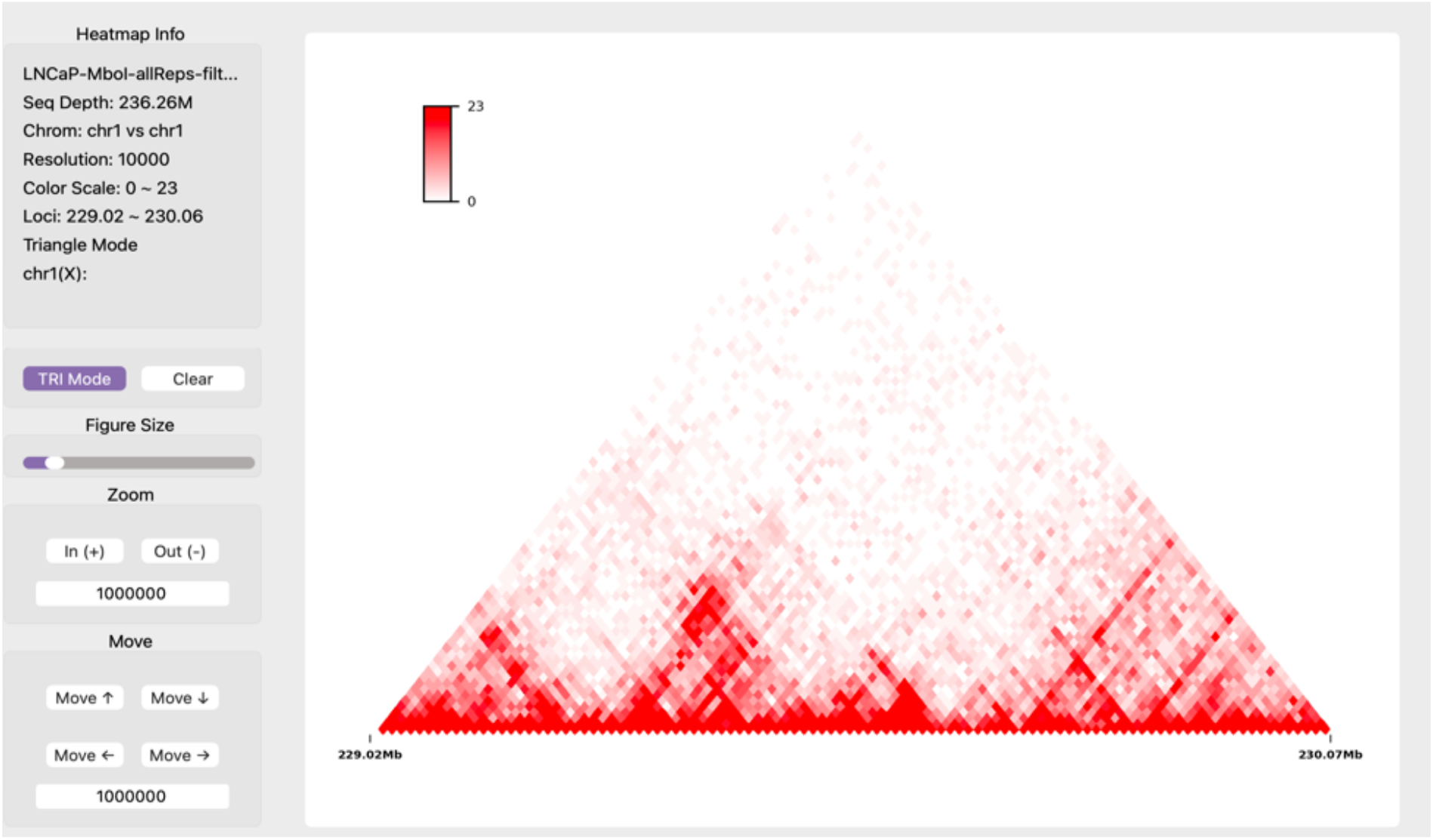

**Tutorial Figure 10.**
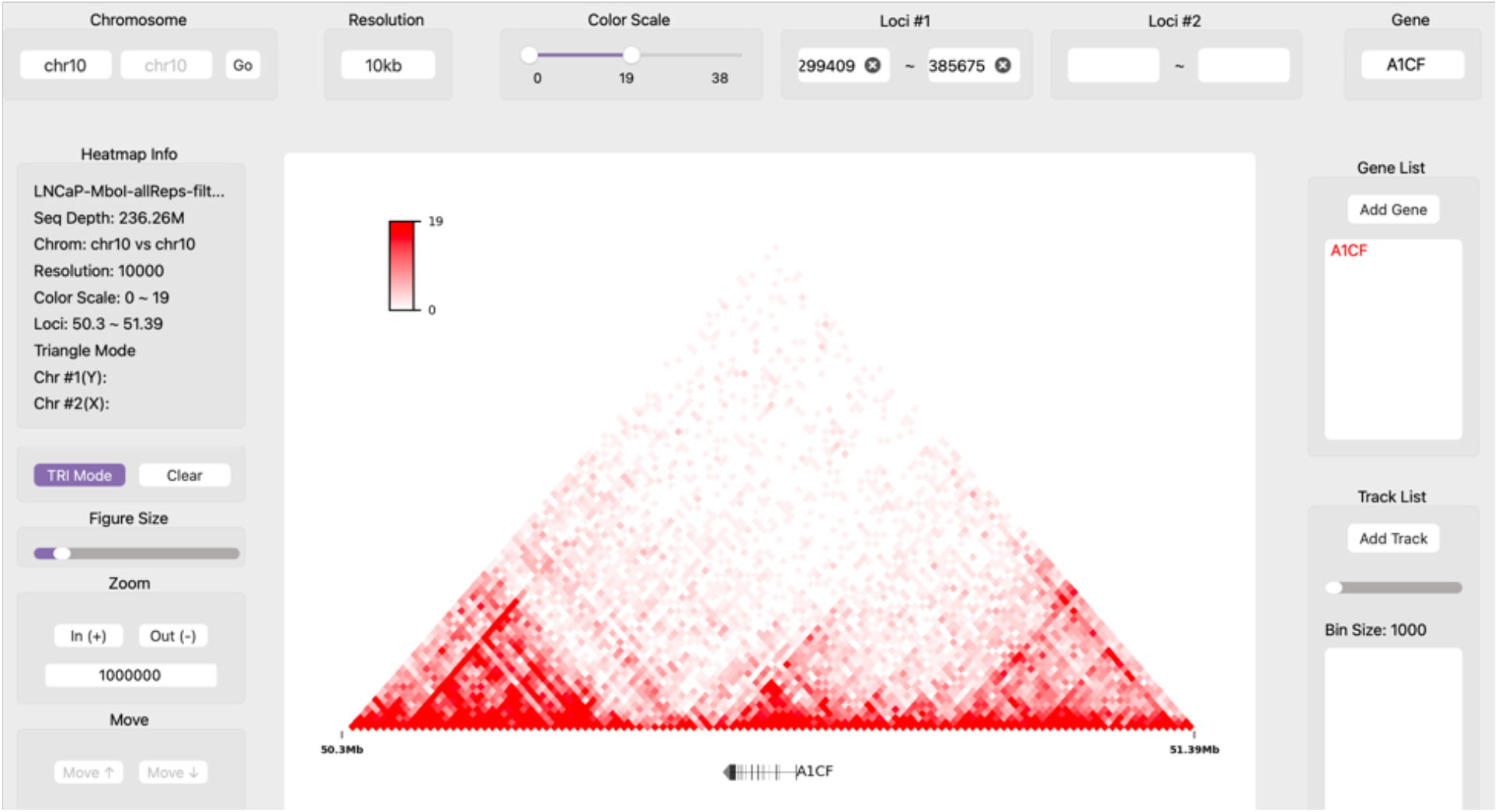

##### 2.5 Navigate Hi-C maps by genes

SV Discovery also supports navigating Hi-C maps by genes through the gene input box on top-right corner of the application. By default, a Hi-C map centered at the transcription start site of the input gene will be displayed, with an extended region size of 5Mb (**Tutorial Figure 10**).

##### 2.6 Select which genes to plot

When adding the gene track, users can select which genes to plot from the drop-down gene list. After clicking on the “Add Gene” button in the right panel, a sub-window will pop up listing all genes in current region (**Tutorial Figure 11**).

**Tutorial Figure 11.**
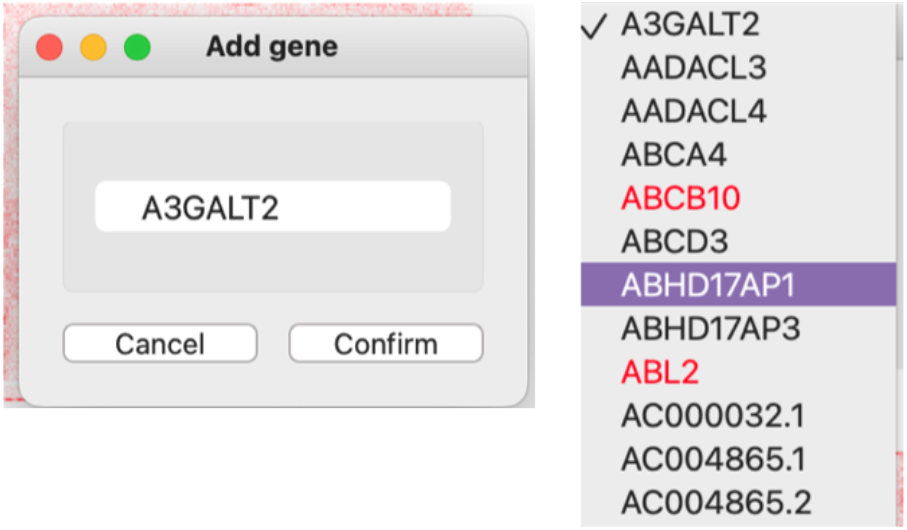

The selected gene names will be listed in the box below the “Add Gene” button. Users can check the information of each gene by either double clicking a gene name in this list, or double clicking a gene in the gene track.

The plotted genes and gene names can be hided any time by using “Gene Display” in the “Settings” menu. User can also remove the gene track by “Clear All Genes” in the “Functions” menu (**Tutorial Figure 12**).

**Tutorial Figure 12.**
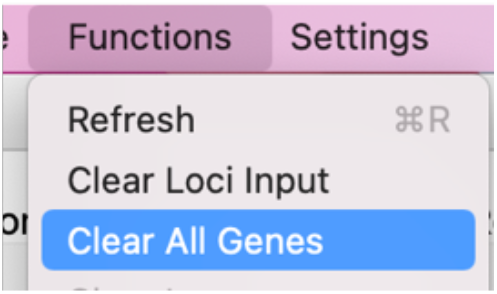

##### 2.7 Add a BigWig track

EagleC Explorer supports plotting one-dimensional (1D) tracks in the BigWig format (https://genome.ucsc.edu/goldenpath/help/bigWig.html). To load a BigWig file, users can either click “import bigWig” from the “File” menu, or click the “Add Track” button. The “Bin Size” bar below the “Add Track” button can be used to smooth the signals.

Double clicking a track from the list below the “Bin Size” bar will pop up a window containing parameters to change the name, the color, the height, and the value range of this track (**Tutorial Figure 13**). And users can also select to hide/show the track label from the “Settings” menu.

**Tutorial Figure 13.**
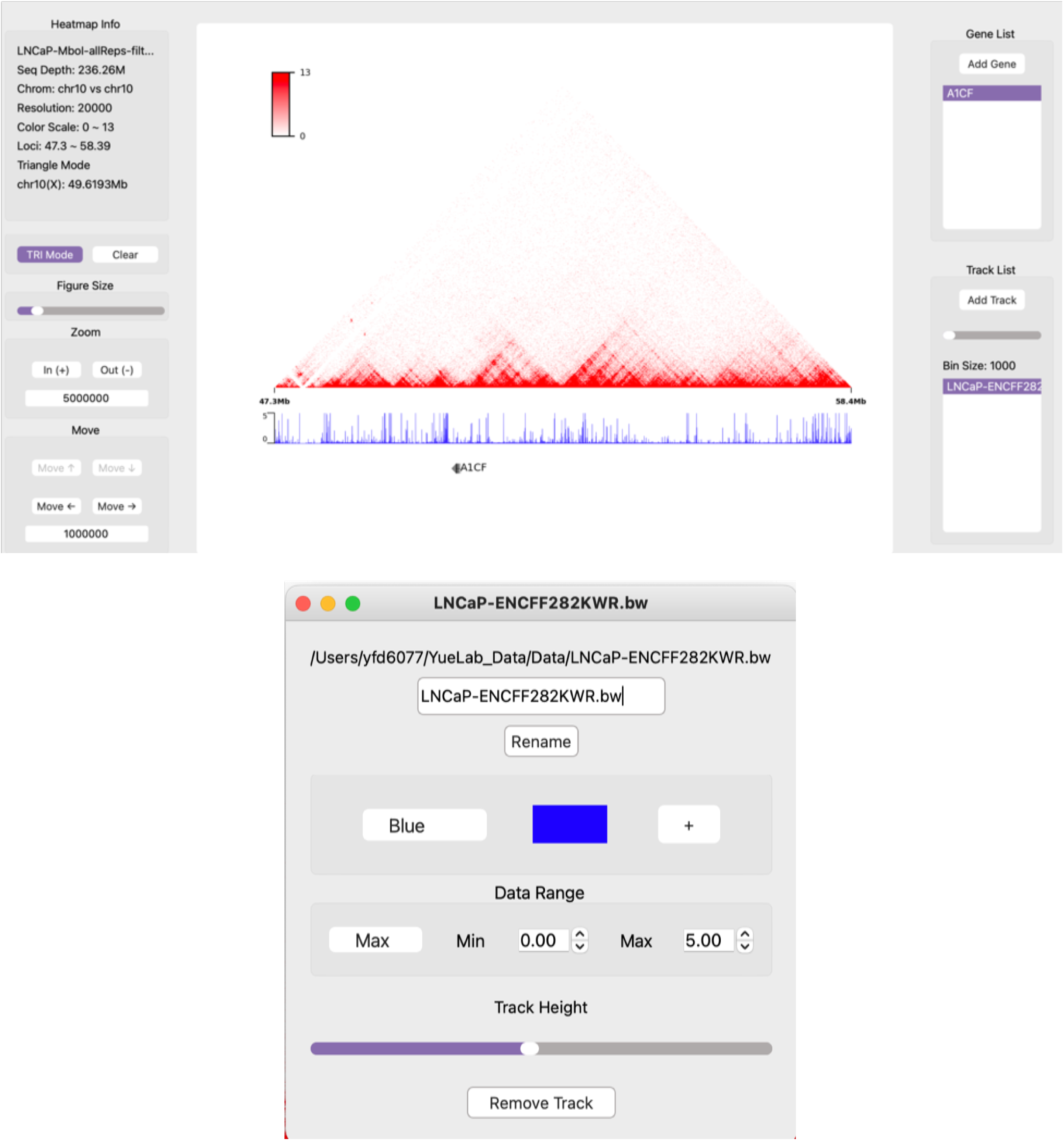

##### 2.8 Plot chromatin loops

EagleC Explorer supports plotting chromatin loops in the BEDPE format (https://bedtools.readthedocs.io/en/latest/content/general-usage.html#bedpe-format). To change the loop color, go to the “Settings” menu and click “Loop Color”. To remove loops, go to the “Functions” menu and click “Clear Loops” (**Tutorial Figure 14**).

**Tutorial Figure 14.**
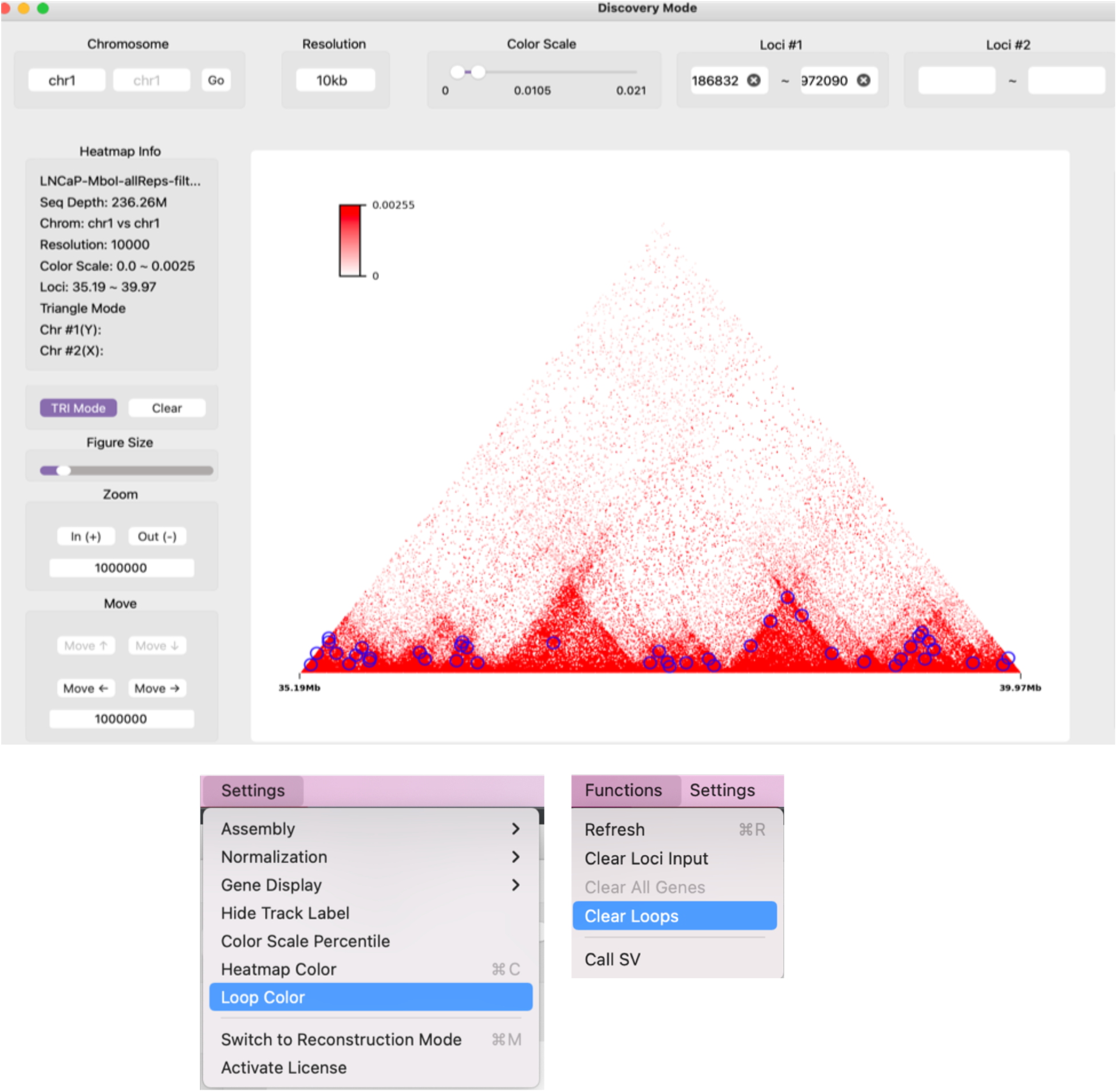

##### 2.9 Zoom in and out of a Hi-C map

EagleC Explorer has three ways for zooming in and out of a Hi-C map: 1) double clicking (or use mouse wheel up) an area on the map will zoom in x3 into the area. 2) Clicking on the “In (+)” and “Out (-)” button will zoom in and out of the map with the specified step size (**Tutorial Figure 15**). 3) drag the mouse cursor over the desired region and click the “zoom in” button in the pop-up dialog.

**Tutorial Figure 15.**
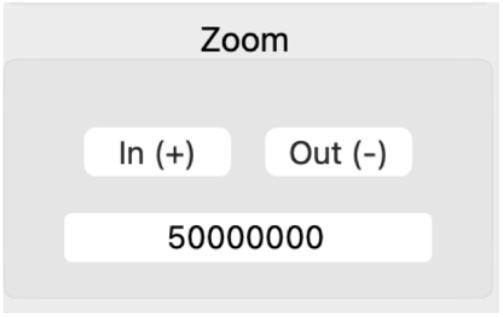

##### 2.10 Move

Users can move a map in 4 directions with a specified step size as well (**Tutorial Figure 16**).

**Tutorial Figure 16.**
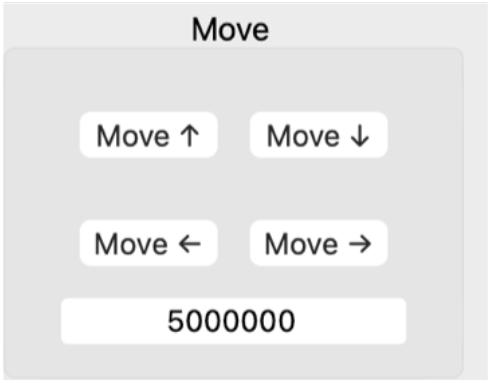

##### 2.11 Interactively detect SVs

As mentioned in 2.9, one way for zooming in a map is to drag the mouse cursor over a desired region and click the “zoom in” button in the pop-up dialog. Clicking on the “call SV” button in the same dialog will invoke the built-in EagleC models to find the precise SV breakpoints within the selected region. After clicking the “Call SV” button, a sub-window will pop up, and users can adjust parameters used for SV calling (**Tutorial Figure 17**). The identified SVs will be exported into the specified file in TXT format under the “EagleC Explorer” folder in users’ home directory.

**Tutorial Figure 17.**
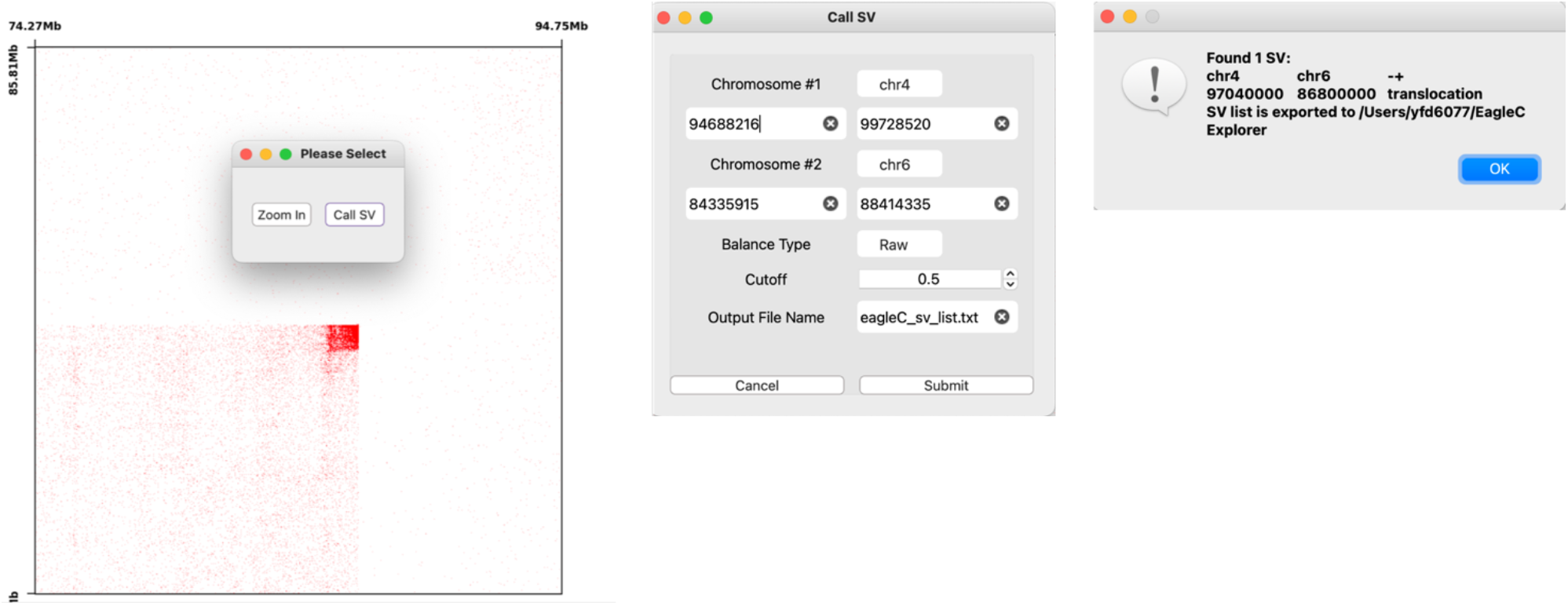

#### 3 The SV Reconstruction Mode

Within the SV Discovery mode, clicking “Switch to Reconstruction Mode” in the “Settings” menu will switch to the SV Reconstruction mode. Most functions in SV Reconstruction are similar to their counterparts in the SV Discovery mode. However, the SV Reconstruction mode is specifically designed for visualizing and reconstructing Hi-C maps surrounding SV breakpoints.

SV Reconstruction generally plots two types of Hi-C maps. On the left canvas, it plots the original Hi-C maps. And on the right canvas, it plots the reconstructed Hi-C maps.

##### 3.1 Load SVs

The top panel of this mode allows users to manually input the breakpoint coordinates of an SV. And the “extend length” determines how far the Hi-C map will be drawn from the breakpoints.

Alternatively, SVs can also be loaded from an external TXT file. The file should contain 6 columns and follow the format described here: https://github.com/XiaoTaoWang/NeoLoopFinder#format-of-the-input-sv-list. Each line in the file corresponds to the information of one SV. And sequentially, the six fields are:

7. Chromosome of the first breakpoint
8. Chromosome of the second breakpoint
9. Orientation type of the SV. Possible values are “+-”, “+-”, “-+”, or “--”.
10. Chromosome position of the first breakpoint
11. Chromosome position of the second breakpoint
12. SV type. Values can be “deletion”, “inversion”, “duplication”, “translocation”, etc.

To load an SV list, just click “Import SV List” in the “File” menu. And once loaded, users can browser them from the top-right drop-down SV list (**Tutorial Figure 18**). Clicking an SV from the list will plot the Hi-C map surrounding that SV breakpoint on the left canvas (**Tutorial Figure 19**).

**Tutorial Figure 18.**
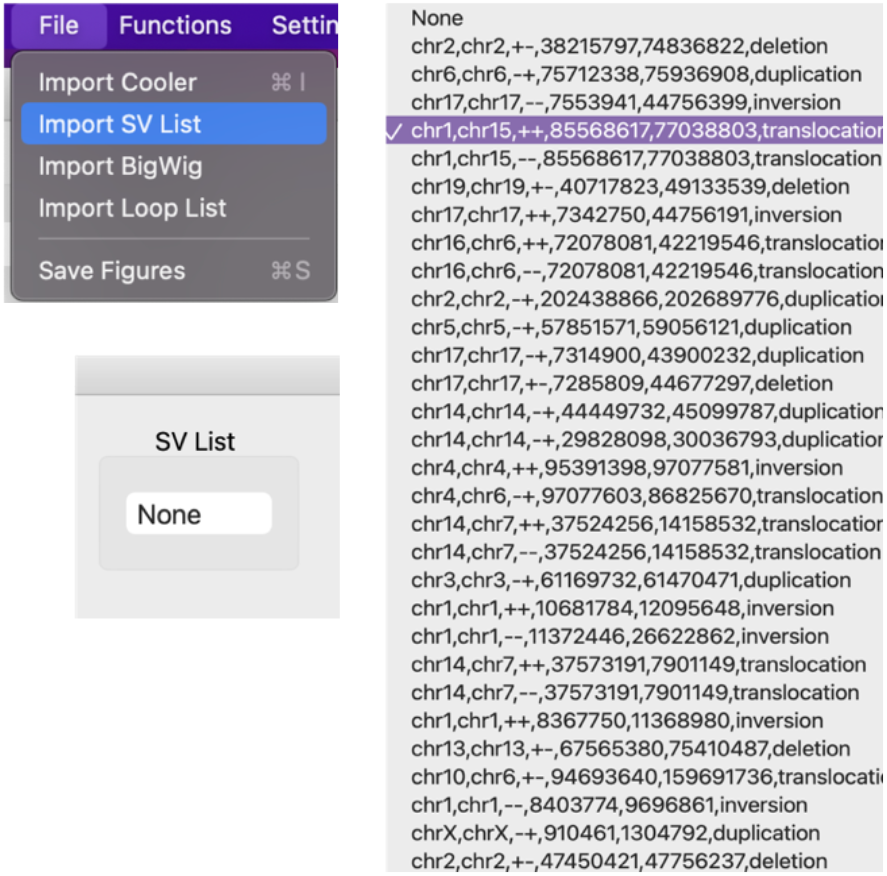

**Tutorial Figure 19.**
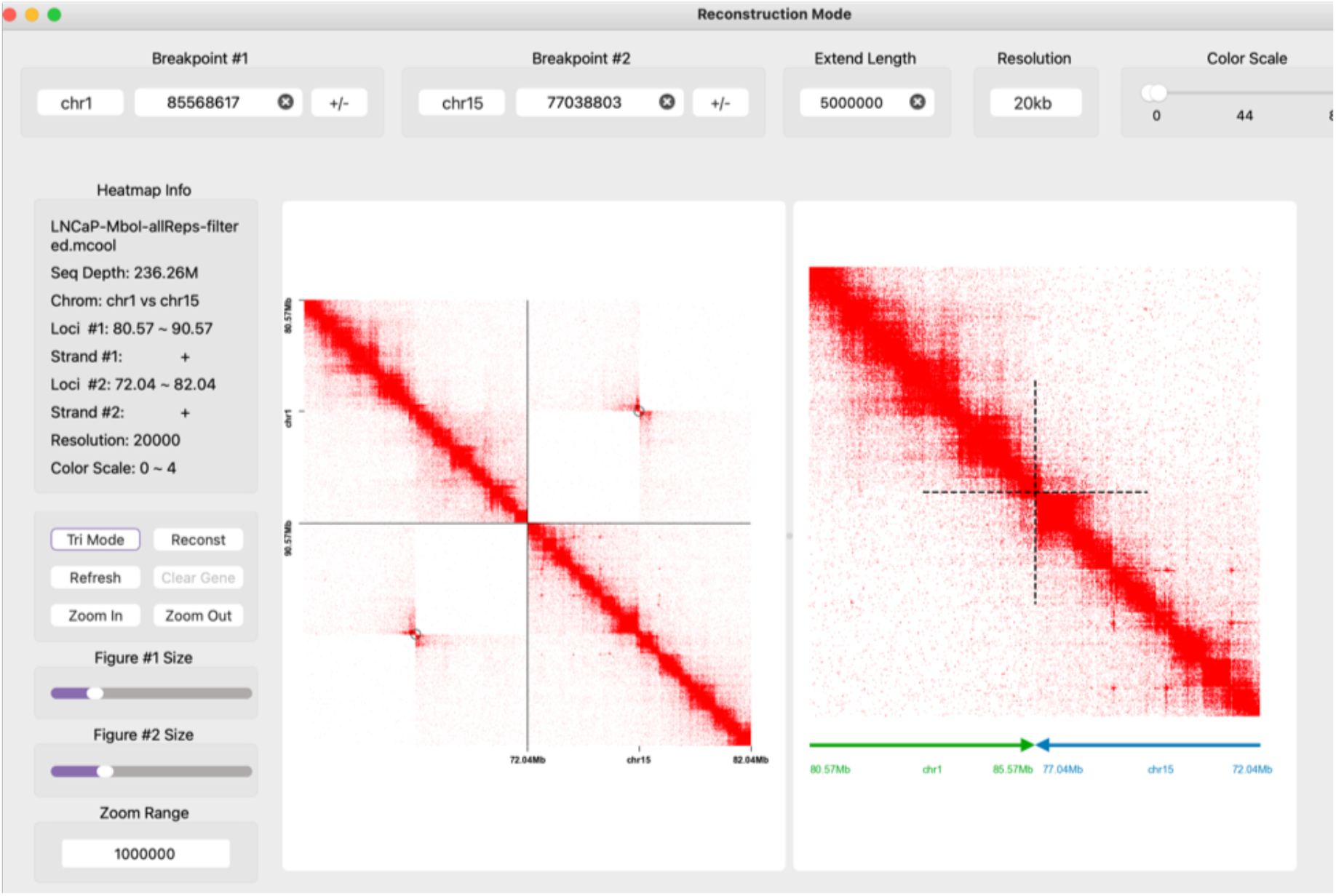

**Tutorial Figure 20.**
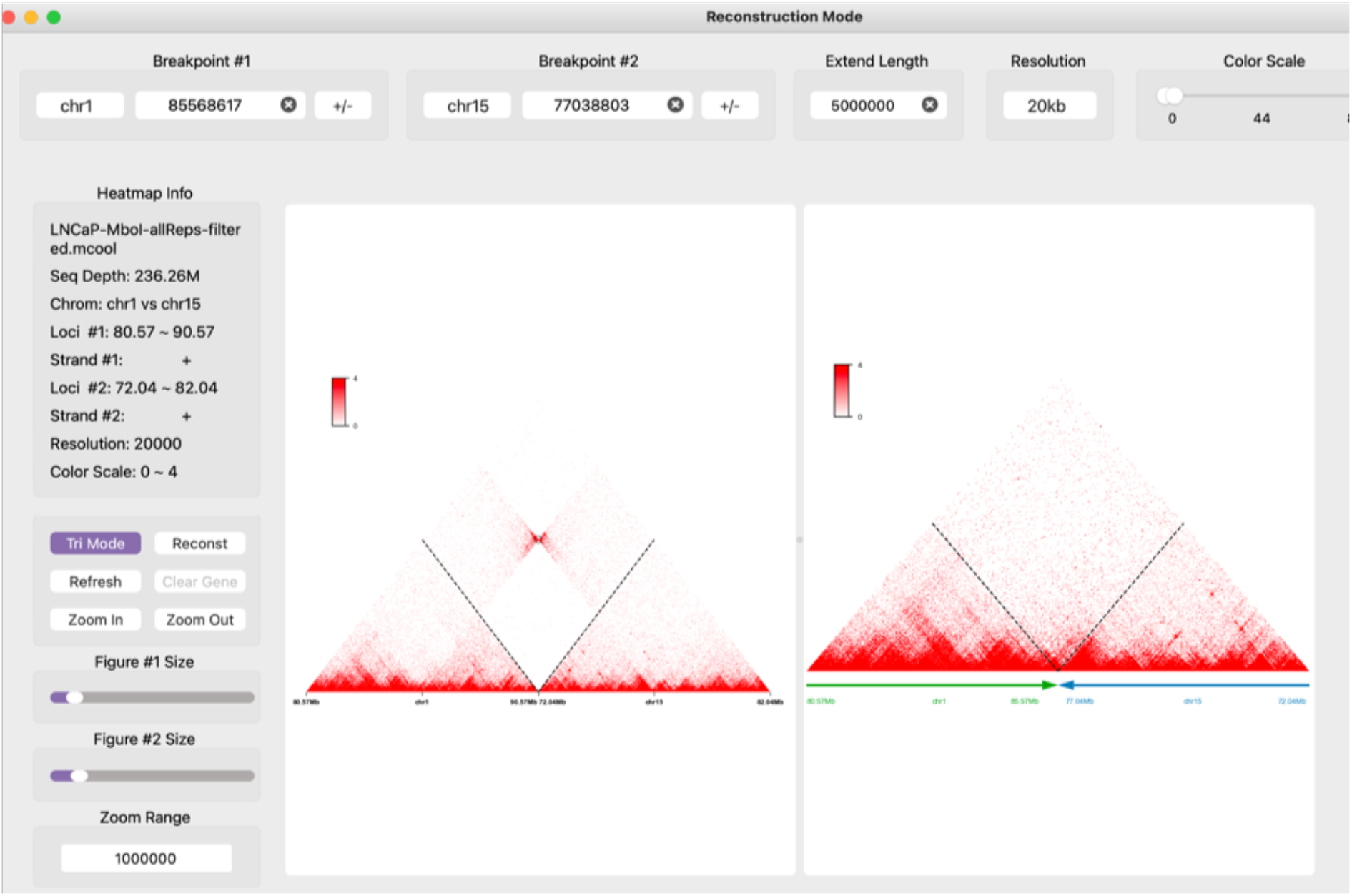

##### 3.2 Reconstruct the Hi-C map

After clicking on the “Reconst” button, EagleC Explorer will reconstruct a local genome with SV fragments placed in the correct order and orientation, and plot Hi-C maps based on this local SV assembly on the right canvas (**Tutorial Figure 19**). Under the heatmap are the chromosome bars, with arrows indicating the fusion orientations. Clicking on the “Tri Mode” button will plot Hi-C maps in a “triangle” format (**Tutorial Figure 20**). By default, both plots will be shown. However, users can drag the boundary of the two canvas to slide out any of them to just focus on one map (**Tutorial Figure 21**).

**Tutorial Figure 21.**
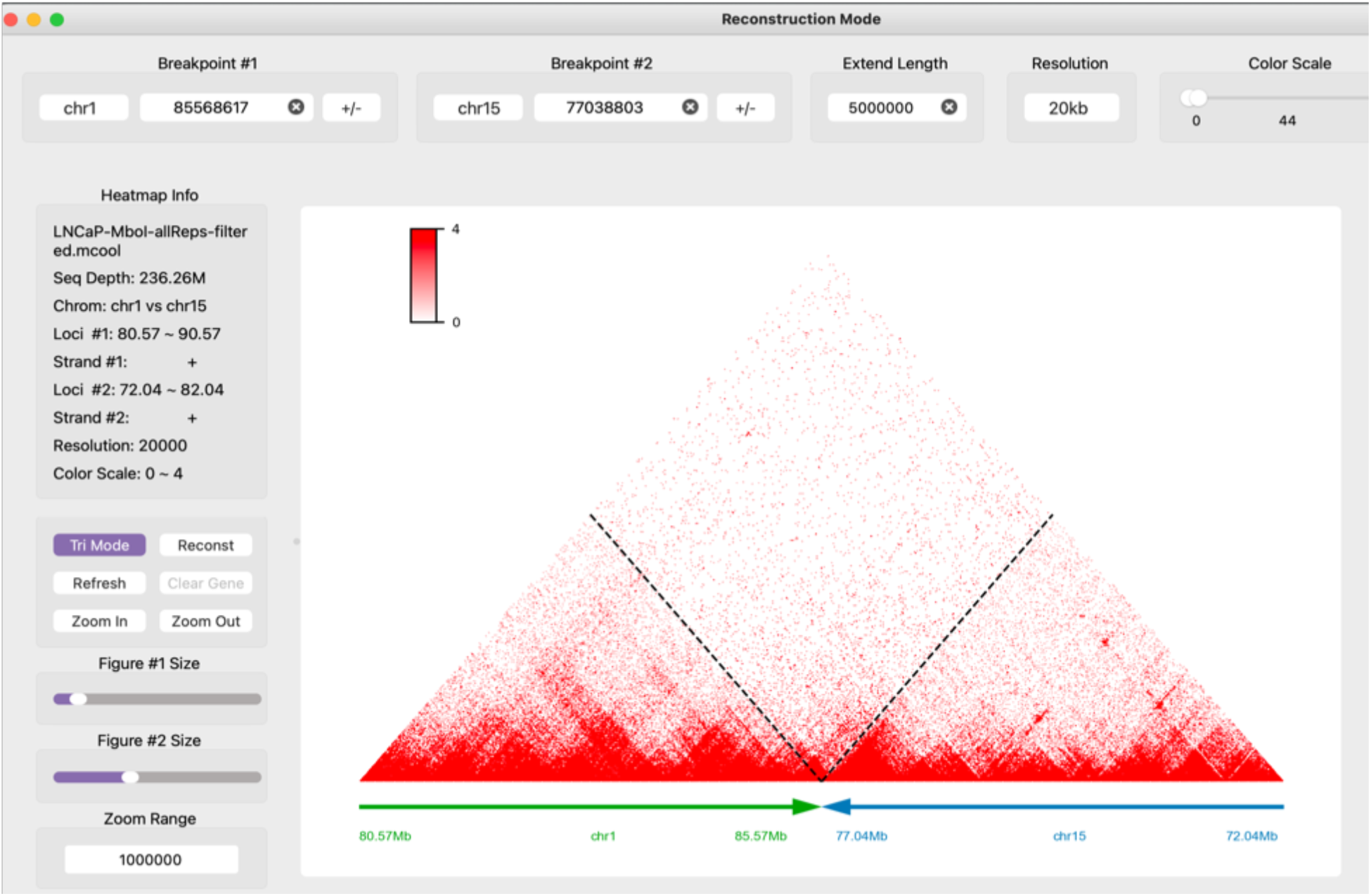

And again, clicking on the “Zoom In” and “Zoom Out” buttons will zoom in and out of the Hi-C maps using the step size specified in the “Zoom Range” Text box.

Users can also add BigWig tracks and plot chromatin loops in a similar way as in the SV Discovery mode. And the sizes of the left canvas and right canvas can be adjusted by using the “Figure #1 Size” and the “Figure #2 Size” bars, respectively.

##### 3.3 Adjust the color scale for interactions across the breakpoints

Due to the heterozygosity of SVs and potential heterogeneity of patient samples, Hi-C signal intensity across the breakpoints tends to be lower than signals in the nearby regions not affected by SVs. EagleC Explorer provides a way to adjust the color scale for signals across the breakpoints alone. To do so, just click the “Interaction Color Scale” in the “Settings” menu, and select a specific coefficient (x1, x1.5, x2, x2.5, x3, x3.5, and x4) you want to multiply on the original signals (**Tutorial Figure 22**).

**Tutorial Figure 22.**
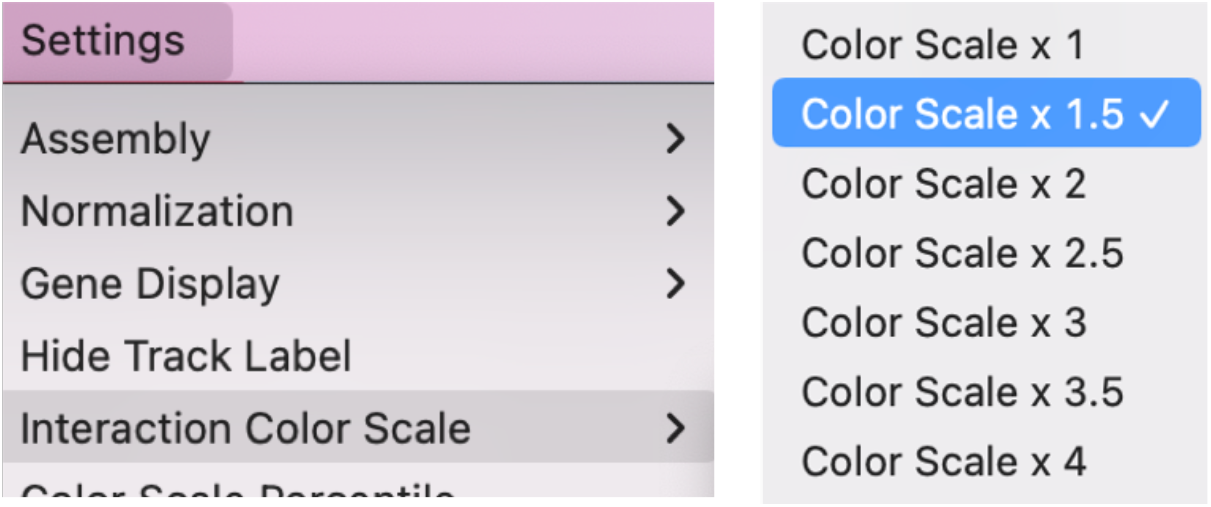

In Tutorial Figure 23, the upper panel shows the original signals, and the lower panel shows 4 times of the original signals.

**Tutorial Figure 23.**
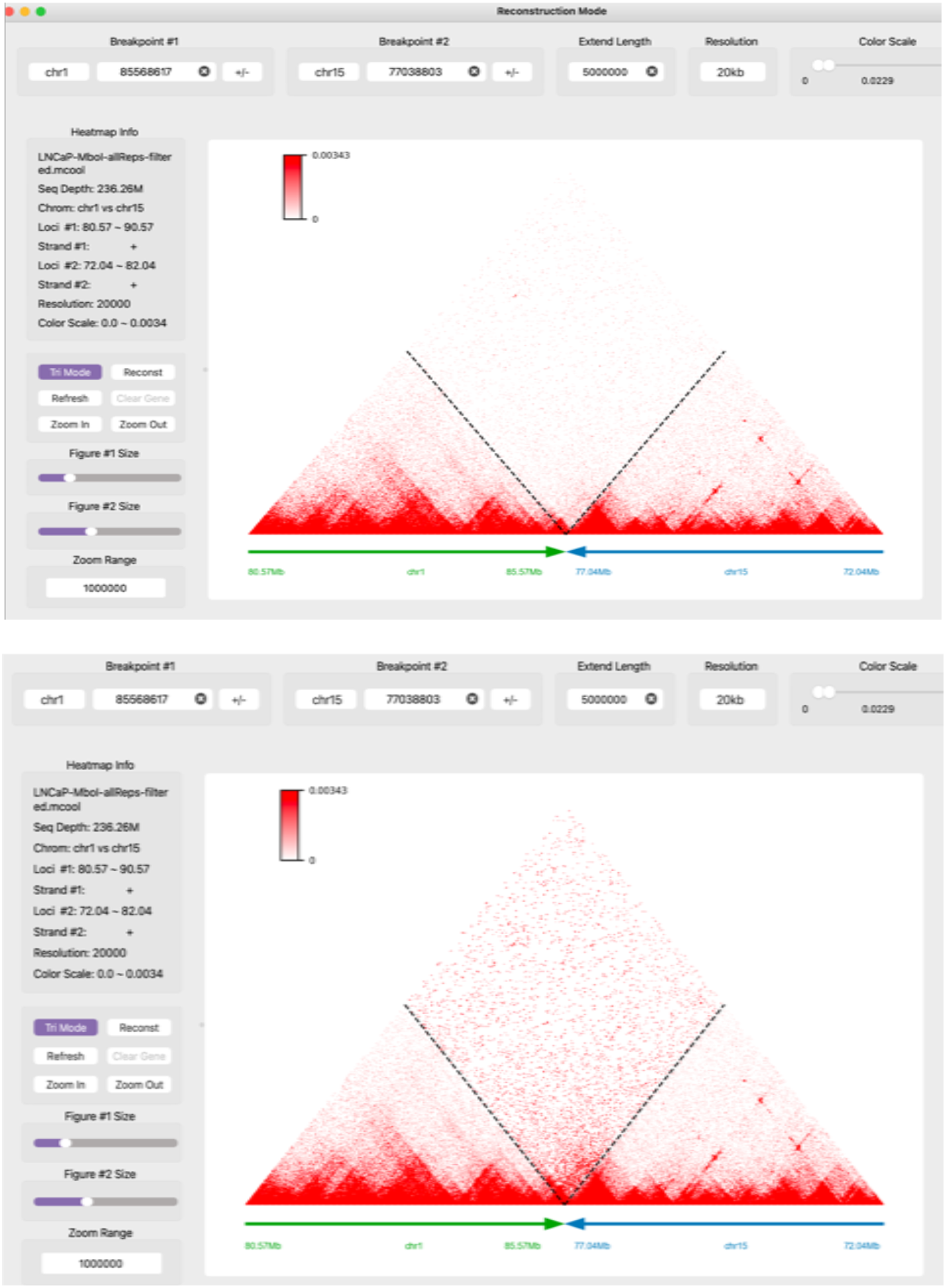

##### 3.4 Add genes

In the SV Reconstruction mode, the “Add Gene” function is specifically customized for better organizing genes surrounding breakpoints. In the drop-down gene list, the genes are divided into two groups, one group are genes surrounding “Break Point #1”, and the other group are genes surrounding “Break Point #2” (**Tutorial Figure 24**). And in both groups, the genes are sorted by the genomic distance to breakpoints and cancer-related genes are highlighted in red (**Tutorial Figure 25**). Clicking on the “Clear Gene” button in the left control panel will remove all genes from the plots. And again, the gene information will be displayed by double clicking a gene.

**Tutorial Figure 24.**
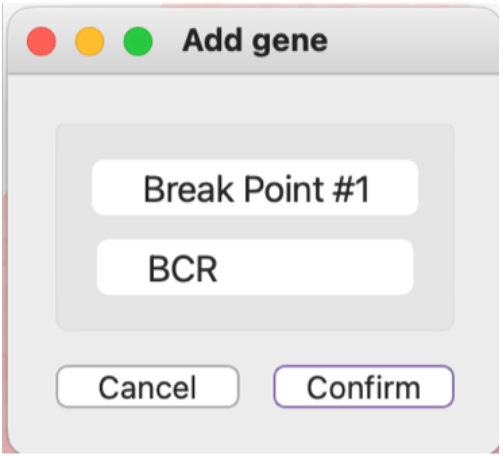

**Tutorial Figure 25.**
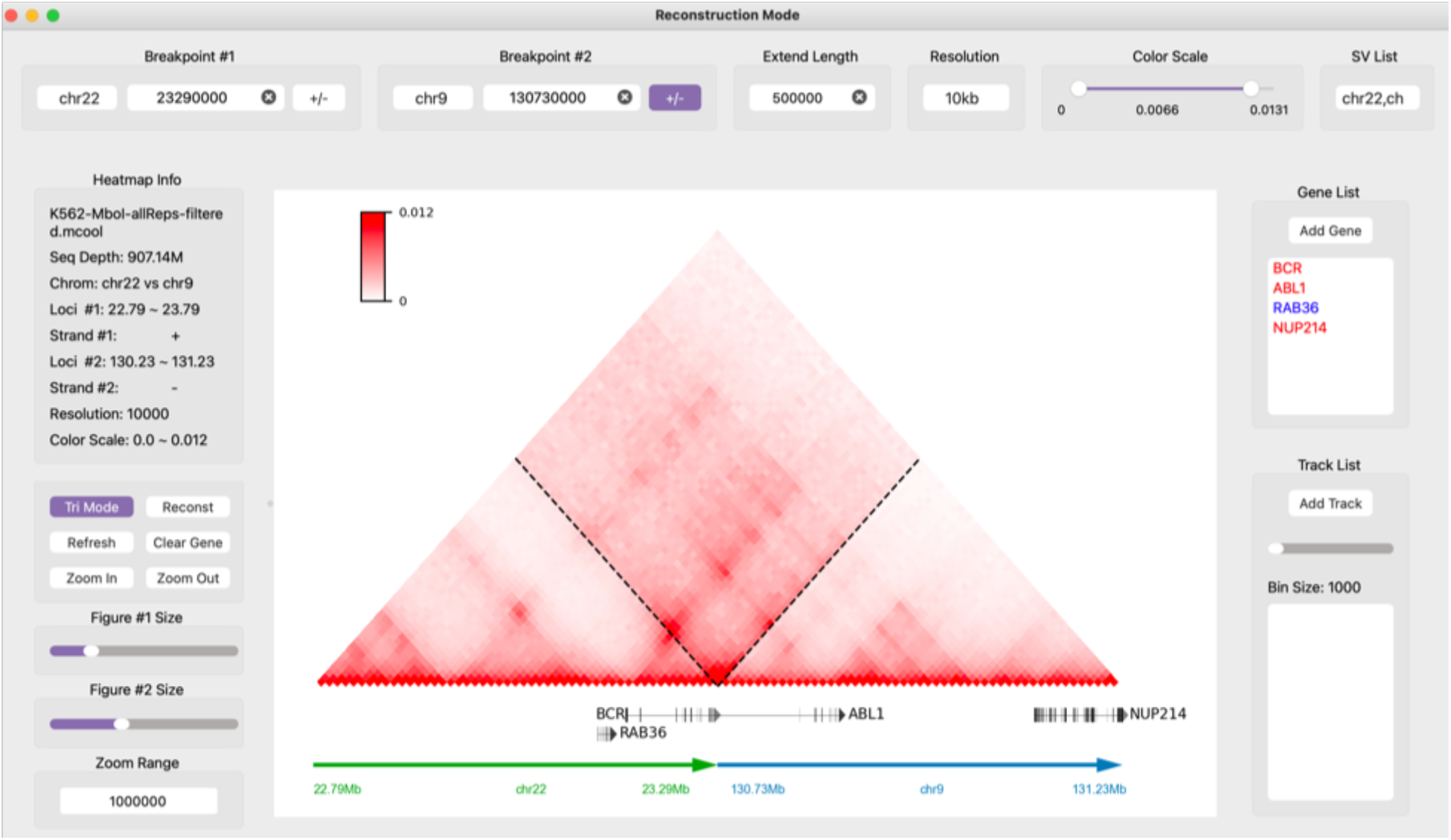

#### 4 Other Functions

##### 4.1 Normalization

EagleC Explorer is able to display Hi-C maps in raw, ICE-normalized, or CNV-normalized values. Users can select the normalization type from the “Settings” menu (**Tutorial Figure 26**). By default, the raw signals will be shown.

**Tutorial Figure 26.**
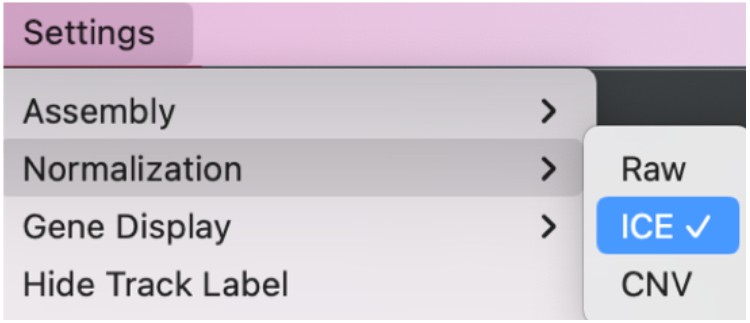

##### 4.2 Heatmap color

The color of a Hi-C map can be changed through the “Heatmap Color” from the “Settings” menu. The default color is red.

##### 4.3 Loop color

The loop color can also be changed from the “Settings” menu. The default color is blue.

##### 4.4 Save figure

The figure can be saved in various image formats through “Save Figure” from the “File” menu. The format of the outputted figure is determined by the suffix of the specified file name.

#### 5 Keyboard Shortcuts

Command + I: Import cooler file

Command + S: Save figure

Command + R: Refresh the canvas

Command + C: Set heat map color

Command + M: Switch between the two modes I: Zoom in

O: Zoom out

+: Increase figure(s) size

-: Decrease figure(s) size

A: Move left (Discovery mode)

W: Move Up (Discovery mode)

S: Move down (Discovery mode)

D: Move right (Discovery mode)

C: Back to the chromosome-wide view (Discovery mode)

P: Draw the original heat map on left canvas (Reconstruction mode)

R: Reconstruct the heat map on right canvas (Reconstruction mode)

